# A FRET assay to quantitate levels of the human β-cardiac myosin interacting heads motif based on its near-atomic resolution structure

**DOI:** 10.1101/2024.12.05.626936

**Authors:** Rama Reddy Goluguri, Piyali Guhathakurta, Neha Nandwani, Aminah Dawood, Seiji Yakota, Osha Roopnarine, David D Thomas, James A. Spudich, Kathleen M. Ruppel

## Abstract

In cardiac muscle, many myosin molecules are in a resting or “OFF” state with their catalytic heads in a folded structure known as the interacting heads motif (IHM). Many mutations in the human β-cardiac myosin gene that cause hypertrophic cardiomyopathy (HCM) are thought to destabilize (decrease the population of) the IHM state. The effects of pathogenic mutations on the IHM structural state are often studied using indirect assays, including a single-ATP turnover assay that detects the super-relaxed (SRX) biochemical state of myosin functionally. Here we develop and use a fluorescence resonance energy transfer (FRET) based sensor for direct quantification of the IHM state in solution. The FRET sensor was able to quantify destabilization of the IHM state in solution, induced by (a) increasing salt concentration, (b) altering proximal S2 tail length, or (c) introducing the HCM mutation P710R, as well as stabilization of the IHM state by introducing a dilated cardiomyopathy-causing mutation (E525K). Our FRET sensor conclusively showed that these perturbations indeed alter the structural IHM state. These results establish that the structural IHM state is one of the structural correlates of the biochemical SRX state in solution.

## Introduction

In heart muscle, force is produced by sliding of myosin-containing thick filaments along actin-containing thin filaments (Geeves, 1991; A. F. Huxley & Niedergerke, 1954; H. E. Huxley, 1965; H. Huxley & Hanson, 1954). Cardiac output is regulated *via* both thin and thick filament-based mechanisms (Brunello et al., 2023; Brunello & Fusi, 2024; Irving, 2024). The latter regulate the availability of myosin heads for interaction with actin and force production (Irving, 2017; Linari et al., 2015). In an actively contracting muscle, not all myosin heads are involved in force production. To conserve energy, and to maintain a reserve from which active heads can be recruited when energy demands increase, some myosin heads in thick filaments exist in a catalytically inactive OFF-state, whose structural basis is the interacting heads motif (IHM) conformation of myosin (Figure 1) (Craig & Padrón, 2022; Lee et al., 2018; Woodhead et al., 2005). In the IHM state, motor activity is shut down by an asymmetric interaction of the two heads (subfragment-1 or S1) of the myosin and folding back of the heads onto the proximal part of their own coiled-coil tail (subfragment-2 or S2) (Grinzato et al., 2023).

**Figure 1.**
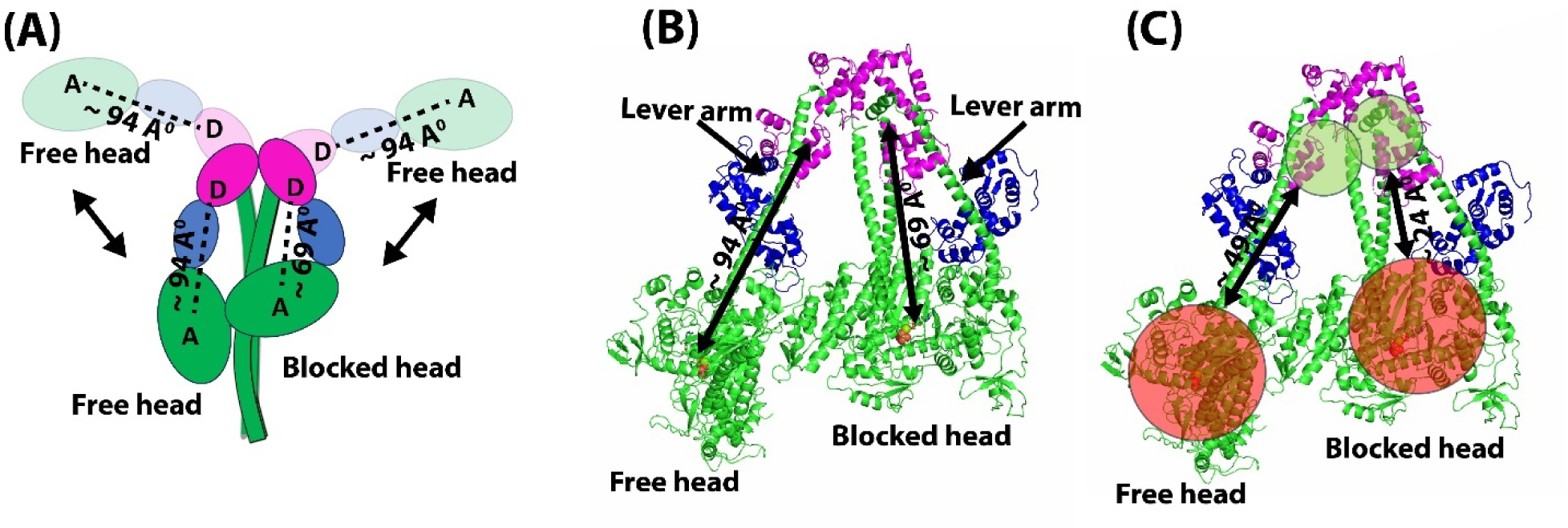
Distances monitored by the AF488 RLC-Cy3ATP or FRET sensor. (A) Schematic diagram showing the open state and the IHM state of β-cardiac HMM. The AF488 donor fluorophore is at the C-terminus of the human ventricular RLC, labeled “D”, and the acceptor Cy3-ATP location is labeled “A”. The distances monitored by the FRET sensor are shown with dashed lines. (B) Near-atomic resolution structure of the IHM conformation of human β-cardiac 15-hep HMM (PDB ID: 8ACT) showing the two distances probed by the FRET sensor. (C) Structure of IHM state showing the distances sampled by linkers of Donor and Acceptor dyes. The range of distances sampled by the donor AF488, which has a ∼15 Å-long C5 linker, are shown in green circles. The range of distances sampled by the acceptor Cy3-ATP, which has a ∼30 Å-long linker, are shown in red circles.

Many HCM-linked β-cardiac myosin mutations that cause hypercontractility of the heart appear to disrupt key S1-S1 or S1-S2 interactions stabilizing the IHM state, subsequently increasing the number of heads available for interaction with actin (Adhikari et al., 2019; Alamo et al., 2017; Barefield et al., 2023; Nag et al., 2017; Robert-Paganin et al., 2018; Schmid & Toepfer, 2021; Spudich, 2015; Trivedi et al., 2017). Twenty HCM-causing mutations in β-cardiac myosin have been analyzed by the long-tail/short-tail actin-activated ATPase ratio (LSAR) assay, which is a direct measure of an increase in myosin heads available for interaction with actin. The LSAR assay compares the effect of mutations on two-headed myosin constructs that either can (long-tail) or cannot (short-tail) form the IHM OFF-state. Of the 20 HCM-mutant myosins analyzed by this direct assay, 19 have been shown to increase the number of heads available for interaction with actin, presumably by destabilizing the IHM state, which has been proposed to be the unifying hypothesis for the mechanism causing HCM-induced hypercontractility of the heart (Spudich et al., 2024).

Dilated cardiomyopathy (DCM) mutations in myosin cause hypocontractility of the heart. One DCM mutation (E525K), in contrast to the HCM mutations, has been reported to stabilize the IHM state, which would result in hypocontractility due to fewer myosin heads being available for force production (Duno-Miranda et al., 2024; Rasicci et al., 2022).

Given the important role played by the IHM state in cardiac regulation, and its association with the pathophysiology of cardiomyopathies, there has been strong interest in structural characterization of the cardiac myosin IHM state, which has been observed in both intact muscle fibers and purified recombinant or native heavy meromyosin (HMM) preparations (Dutta et al., 2023; Grinzato et al., 2023; Tamborrini et al., 2023; Zoghbi et al., 2008). IHM in intact muscle fibers has been probed by electron microscopy (Craig et al., 1992; Kensler, 2002), X-ray diffraction (Ma & Irving, 2022; Matsubara & Millman, 1974), fluorescence polarization (Brunello et al., 2023; Linari et al., 2015), and cryo-electron microscopy (Dutta et al., 2023; Tamborrini et al., 2023). The latter has recently revealed exquisite details of the structure of the thick filaments under relaxed conditions (Dutta et al., 2023; Tamborrini et al., 2023). These techniques, however, do not allow a systematic analysis of the molecular effects of the hundreds of disease-causing mutations in cardiac myosin. Recombinant human β-cardiac myosin protein expressed and purified from differentiated mouse myoblast cells has proved to be an excellent alternative (Resnicow et al., 2010; Sommese et al., 2013; Srikakulam & Winkelmann, 2004) that has been widely used to understand the effects of disease-linked mutations on myosin’s structure and functions (Spudich et al., 2024).

Several assays have been developed to study transitions between ON and OFF states of purified β-cardiac myosin protein, the most direct of which is the LSAR assay described above. Another biochemical assay that often has been assumed to be a quantitative measure of the IHM OFF-state is a single nucleotide turnover assay that reports on the energy-conserving SRX state of myosin (Cooke, 2011; Craig & Padrón, 2022; Hooijman et al., 2011; Nag & Trivedi, 2021; Stewart et al., 2010). The SRX assay has been used for purified protein preparations as well as for fibers and cells. In the SRX assay, myosin binds and hydrolyzes the fluorescent ATP analog 2’/3’-O-(N-Methyl-anthraniloyl)-adenosine-5’-triphosphate (Mant-ATP), and the displacement of Mant-ADP by dark ATP results in a multi-exponential fluorescence decay arising from different enzymatic states of myosin with distinct ATP turnover rates. The IHM OFF-state is thought to correlate with SRX myosin, which has an extremely slow rate of ATP turnover, typically ∼1000-fold slower than the actin-activated ATPase rate and ∼10-fold slower than the basal rate of ON-state myosin heads when they are not bound to actin (disordered relaxed state or DRX). Many HCM-causing mutations in cardiac myosin have been found to destabilize the SRX, several of which have been independently shown to weaken S1-S2 binding, which is part of the protein-protein interactions that stabilize the IHM (Adhikari et al., 2016; Nag et al., 2017). In contrast, the small molecule myosin inhibitor mavacamten (Camzyos), which is used to treat obstructive HCM (Green et al., 2016; Nag et al., 2023; Owens et al., 2024), and two DCM mutations in myosin, which cause hypocontractility of the heart, have been shown to stabilize the SRX state (E525K in myosin: Duno-Miranda et al., 2024 and Rasicci et al., 2022; P94A in the regulatory light chain (RLC) of myosin: Yuan et al., 2018). In most cases, there is qualitative agreement between the biochemical SRX state and the IHM OFF state, as indicated by the general correlation between results from the SRX assay, the LSAR assay, negative staining EM, and fiber X-ray diffraction, all of which report on the availability of myosin heads (Spudich et al., 2024).

There are, however, several major issues in quantifying the IHM state indirectly from the SRX assay. First, additional structural states of myosin other than the IHM have been shown to result in slow turnover of ATP (Chu et al., 2021; Craig & Padrón, 2022). Single-headed myosin constructs and double-headed myosin constructs with a short tail that cannot form the IHM still show 10-20% SRX (Anderson et al., 2018; Duno-Miranda et al., 2024; Rohde et al., 2018). The structural origin of the SRX state in these constructs is unknown. Additionally, in the presence of myosin modulators, a discrepancy is observed between SRX measurements and structural studies measuring the ordered arrangement of myosin heads (X-ray diffraction). Both omecamtiv mecarbil (OM) and piperine have been shown to disrupt the ordered arrangement of myosin heads without altering the SRX fraction in muscle preparations (Jani et al., 2024). Similarly, the myosin activator 2’-deoxy-ATP (dATP) has been shown to completely disrupt the biochemical SRX state, but the ordered arrangement was disrupted only by 25% (Ma et al., 2023). Finally, there are 6 examples out of 20 HCM mutants studied where the correlation between the SRX state and the number of myosin heads available to form cross bridges with actin, as measured by the LSAR assay, is lost (Spudich et al., 2024). In these cases, the nucleotide turnover assay shows no change in the SRX/DRX ratio of the mutant as compared to WT, while the LSAR assay reveals that the mutation results in more heads available to interact with actin (Spudich et al., 2024). So, caution must be used in interpreting the SRX assay results alone for quantitating IHM state.

Both the LSAR assay and the nucleotide turnover assay are biochemical measurements, so an assay that monitors the IHM structural state directly is highly desirable. Fluorescence resonance energy transfer (FRET) is an obvious choice to examine the structural changes from (a) open myosin heads free to interact with actin to (b) folded-back heads in the IHM state. Here, we report a FRET-based approach to quantitate IHM state levels in solution, and we use this FRET assay to demonstrate disruption of the IHM state by an HCM-causing mutation and stabilization of the IHM state by a DCM-causing mutation.

## Materials and Methods

### Buffers and reagents

All experiments were performed in 10 mM Tris pH 7.5, 4 mM MgCl_2_, 1 mM EDTA, 1 mM DTT and 0.02% Tween 20 with appropriate amounts of salt (potassium acetate) added for the experiments performed at different salt concentrations. All the buffers were filtered through a 0.22 μm filter. Alexa Fluor^®^ 488 C5 Maleimide (AF488) (Catalog # A10254) was purchased from Thermo Fisher Scientific, and a 10 mM stock of the dye was made in DMSO and stored at −20°C. A 1 mM stock of Cy3 ATP (Catalog # NU-808-CY3) was purchased from Jena Biosciences.

### Expression and purification of human ventricular RLC with C-terminal cysteine from inclusion bodies

Human ventricular RLC (MYL2) with a cysteine residue added at the C-terminus after Asp166 (last amino acid in RLC) was used in the current study (referred to as Cys-RLC henceforth). DNA encoding Cys-RLC was synthesized and subcloned into the pET28b expression vector by GenScript (Piscataway, NJ). This expression plasmid also contains a 6-His tag followed by a TEV protease cleavage site on the N-terminus of the Cys-RLC. The plasmid was transformed into the *E.coli* Rosetta (DE3) pLysS strain for expression and purification. A starter culture was prepared by inoculating a single colony of the transformed cells in 100 mL LB medium containing 30 ug/mL kanamycin. The starter culture was grown overnight (12-16 hrs) at 37°C and 200 rpm shaking and transferred to 2L LB medium containing 30 μg/mL kanamycin. The 2L culture was grown at 37°C and 200 rpm shaking until an OD of 0.6 at 600 nm was achieved. The culture was induced with 1 mM IPTG and grown for another 3 hours. The cells were harvested by centrifuging at 6000 rpm for 20 minutes at 4°C. The cell pellet was dissolved in 1X PBS and centrifuged at 5000 rpm for 10 minutes on a bench top centrifuge to pellet down the cells. PBS was aspirated and cell pellets were flash frozen and stored at −80°C. For protein purification, the cell pellet was thawed on ice, dissolved in a buffer containing 20 mM Tris pH 7.5, 6 M Urea, 1 M NaCl, 1 mM PMSF and 5 mM BME and sonicated in a Vibronics sonicater at 70% Amplitude, 50% duty cycle for 30 seconds each 4 times. The resulting cell lysate was centrifuged at 35,000 rpm for 40 minutes at 4°C. The supernatant was incubated with 5 mL of Ni-NTA beads on a rocker for 1 hour at 4°C. The remaining steps were performed in a 50 mL falcon tube in a batch process mode at 4°C. The resin was washed 2 times with ∼10 column volumes of buffer containing 20 mM Tris pH 7.5, 1M NaCl and 6M urea and then 3 times with buffer containing 20 mM Tris pH 7.5 and 300 mM NaCl. The protein was eluted using buffer containing 20 mM Tris pH 7.5, 300 mM imidazole and 300 mM NaCl. The imidazole was removed by dialyzing the protein solution against 20 mM Tris pH 7.5 and 150 mM NaCl to a million-fold dilution using 3.5 KDa cutoff dialysis tubing (Snakeskin, Cat# 88244, Thermo Fisher Scientific). The purified protein was flash frozen in liquid nitrogen and stored at −80°C.

### Labeling of human ventricular Cys-RLC with AF488

The single cysteine at the C-terminus of the RLC was labeled with AF488. Cys-RLC at 90 µM concentration in 10 mM Tris pH 7.5, 150 mM NaCl, and 8 M urea was incubated with 180 µM AF488 solution for 6 hours at 23°C. Excess dye was removed by performing affinity purification of labeled protein using Ni-NTA resin as described above. The labeled protein was flash frozen in liquid nitrogen and stored at - 80°C. The extent of labeling was found to be 100% by mass spectrometry (Figure S1).

### Purification of human β-cardiac myosin

All constructs of β-cardiac myosin were recombinantly expressed and purified from the mouse myoblast C2C12 cell line. The 8-hep and 15-hep myosin constructs are the same as those used in a previous study(Nandwani et al., 2024). The 15-hep myosin construct contains residues 1-942 of MYH7, which comprises the S1 motor head along with 15 heptad repeats of the proximal S2 region, followed by a GCN4 leucine zipper sequence to ensure dimerization and a GSG linker followed by an octapeptide sequence (RGSIDTWV) that can bind a PDZ domain. The 8-hep myosin construct is identical except that it contains residues 1-893 of MYH7 and includes only the first 8 heptads of the proximal S2 tail. C2C12 cells were differentiated by placing in starvation media for 48 hours and then infected with two adenoviruses carrying the MYH7 gene and the human ventricular ELC (MYL3) gene containing an N-terminal FLAG tag followed by a TEV cleavage site. The infected cells were harvested after 4 days, and the cell pellet was flash frozen and stored at −80°C. For protein purification, the cell pellet was thawed, and the recombinant human β-cardiac myosin was purified by affinity chromatography using an anti-FLAG resin as described previously(Morck et al., 2022; Nandwani et al., 2024). The endogenous mouse RLC was completely depleted (Supplementary Figure S2) and exchanged with the AF488 labeled Cys-RLC while the protein was still bound to the anti-FLAG resin, as described previously (Morck et al., 2022; Nandwani et al., 2024). The resin was washed extensively after the binding step to completely remove free dye. The resultant myosin, containing only AF488 labeled RLC, was eluted from the anti-FLAG resin by treating it with TEV protease overnight. The protein was further purified by anion exchange chromatography using a 1 mL Q Sepharose FF column on an AKTA FPLC system. The fractions containing pure human β-cardiac myosin without endogenous mouse myosin were pooled and flash-frozen in liquid nitrogen and stored at −80°C.

### Steady state FRET measurements

Steady-state FRET measurements were performed in an Infinite M Nano+ fluorescence plate reader (Tecan). β-cardiac myosin containing AF488 labeled Cys-RLC was buffer exchanged into assay buffer using an Amicon Ultra Centrifugal filter device with 50 KDa molecular weight cut off. FRET measurements were performed in a buffer containing 10 mM Tris pH 7.5, 4 mM MgCl_2_, 1 mM EDTA, 1 mM DTT and 0.02% Tween 20. The salt titration was performed by adding appropriate amounts of potassium acetate salt to the buffer. 10 nM labeled protein was mixed with 1 µM Cy3 ATP and incubated for 5 minutes to reach equilibrium. The fluorescence emission spectra of AF488-labeled Cys-RLC of human β-cardiac myosin was obtained by exciting at 470 nm, and the emission spectrum was recorded from 500 nm to 600 nm with 1 nm increments and a gain value of 100. The signal integration time was 2 ms. Blank spectra were obtained from the sample containing 1µM Cy3 ATP without any protein. All the spectra were blank subtracted, and FRET efficiency (E) was calculated from the blank subtracted emission spectra using the following formula:

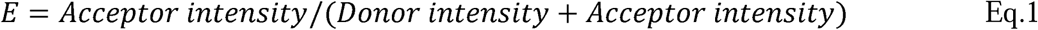

### Time-resolved FRET (TR-FRET) experiments

Time-resolved fluorescence measurements were performed using a custom-built spectrophotometer equipped with a time-correlated single-photon counting card SPC 180 N (Becker and Hickl GmbH, Berlin). The time-resolved fluorescence waveforms were recorded after exciting the AF488 donor fluorophore at 470 nm using a picosecond pulsed laser source LDH-I (PicoQuant GmbH, Berlin) pulsing at 60 MHz. The fluorescence from the sample was filtered through a band pass filter placed at magic angle and detected using a PMH-100 photomultiplier tube (Becker and Hickl GmbH, Berlin). The count rate from the sample was adjusted to 0.5-0.6 MHz to prevent pulse pileup. Each sample was acquired 25 times and averaged to generate a final waveform. The instrument response function (IRF) was obtained by collecting the scattered light from a water sample in the cuvette.

Lifetime decay curves were analyzed by fitting to a multiexponential decay equation,

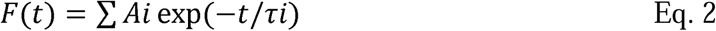

where A represents the amplitude of each phase and τ represents the lifetime. The donor lifetime decay curves were found to best fit to a three-exponential decay equation for both donor-only and donor-acceptor samples. The amplitude-weighted average lifetime was used to calculate FRET efficiency (E) using the equation,

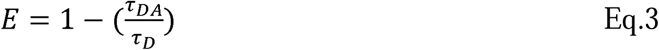

where τ_DA_ is the average donor lifetime in the presence of acceptor (obtained by mixing 20 nM AF 488 labeled myosin with 2 μM Cy3 ATP), and is the average donor-only lifetime (obtained from the sample containing 20 nM AF488-labeled myosin along with 2 µM dark ATP (non-fluorescent).

### Single turnover SRX assay

The single-nucleotide turnover SRX assay for β-cardiac myosin was performed by manual mixing using the acquisition system of the Biologics stopped flow system (SFM 3000). 100 nM myosin was mixed with 200 nM Mant-ATP and incubated for 20 seconds for the binding and hydrolysis of Mant-ATP to occur. The turnover kinetics of Mant-ATP was probed by chasing the reaction with 1 mM unlabeled ATP. The fluorescence of Mant was excited by exciting the tryptophan on myosin at 290 nm which is known to transfer energy to the bound Mant-ATP through FRET. The emission of Mant was observed at 450 nm using a bandpass filter placed in front of the photomultiplier tube. The resulting fluorescence decay was fitted to a double-exponential decay equation to obtain rate constants and amplitudes of the DRX and SRX phases (Anderson et al., 2018).

## Results

### Design of the FRET sensor

The strong distance dependence of FRET was exploited to probe the equilibrium between myosin molecules with open heads and those in the IHM structure with folded back heads. We designed a FRET sensor based on the high-resolution structure of the human β-cardiac myosin IHM (PDB ID: 8ACT) (Grinzato et al., 2023). In cardiac IHM, the lever arm of the blocked head (BH) has a kink that brings the motor domain of the head (labeled green in Figure 1) closer to the lever arm region compared to the free head (FH). We utilized this structural transition in designing the FRET sensor, where the donor fluorophore (AF488) is attached to the C-terminus of the Cys-RLC and the acceptor fluorophore (Cy-3 ATP) is a fluorescent ATP occupying the nucleotide binding pocket (Figure 1). We used the 15-hep heavy meromyosin (HMM) construct of human β-cardiac myosin, which is a hexamer comprised of two S1 heads with essential and regulatory light chains, ELC and RLC, respectively, bound to each lever arm, followed by 15 heptads of the proximal S2 coiled-coil tail (Figure 1). The human β-cardiac 15-hep HMM has been shown to adopt the IHM conformation in solution and its high-resolution structure was recently determined using cryo-EM (PDB ID: 8ACT) (Grinzato et al., 2023; Nandwani et al., 2024). In the cardiac myosin IHM, the motor domains of the two heads adopt the classical pre-powerstroke (PPS) conformation and are essentially superimposable, while the two lever arms are positioned differently, giving rise to the asymmetry in the structure. In the open ON-state there is no kink in the BH pliant region, and the asymmetry is lost. We made use of this structural feature and placed the donor and acceptor fluorophores such that the FRET sensor reliably distinguishes between the open ON-state and the closed IHM OFF-state in solution. We covalently attached a donor (D) fluorophore, AF488, to an engineered cysteine residue at the C-terminus of the human ventricular RLC. Bacterially expressed human ventricular Cys-RLC was purified, fluorescently labeled, and exchanged onto the 15-hep HMM expressed in C2C12 cells during the purification process, as described in Materials and Methods. Fluorescently tagged 15-hep HMM showed the same %SRX as that of the unlabeled 15-hep HMM, indicating that the AF488 tag did not affect the biochemical SRX state (Figure S3). Cy3-ATP bound at the nucleotide binding pocket of the myosin head served as the acceptor (A) fluorophore.

In the cardiac myosin IHM, the distance between the C-terminus of the RLC and the hydroxyl group of the ribose ring of the ADP bound at the nucleotide binding pocket is ∼ 94 Å in the FH and ∼ 69 Å in the BH (Figure 1A and 1B). These distances are approximate, since the disordered C-terminal four residues of the RLC are not fully resolved in the cryo-EM structure; an extra Cys residue has been added to the C-terminus in our construct. The Cy3 and AF488 moieties are large enough to reduce these distances if they were pointing toward one another in the IHM structure or increase the distances if pointing away from one another. Thus, the FRET dynamic range for BH AF488-Cy3ATP distance could be as small as 24 Å or as large as 114 Å, and for the FH AF488-Cy3ATP distance could be as small as 49 Å or as large as 139 Å (Figure 1C) (see Supplemental Figure S4 and Supplementary Methods). The R_0_ value for the AF488-Cy3 FRET pair is ∼68 Å, which is appropriate to resolve these FRET pair distances.

### Salt disrupts the IHM state

Earlier studies from our lab and others have shown that the addition of salt disrupts the IHM state, due to disruption of the electrostatic interactions that stabilize the head-head and head-tail interactions (Anderson et al., 2018; Duno-Miranda et al., 2024; Nandwani et al., 2024; Rasicci et al., 2022; Rohde et al., 2018). We probed the effect of changing salt (potassium acetate, KOAc) concentrations on the IHM structure directly, using the FRET sensor, by steady-state FRET measurements. Representative spectra at 0 and 150 mM KOAc, from which FRET efficiencies were calculated, are shown in Figure 2A. The FRET efficiency decreased with increasing salt concentration (Figure 2B), showing more directly that higher salt destabilizes the IHM structure, resulting in more myosin heads in the open conformation. We also quantified the biochemical SRX state on the same preparations by performing the SRX assay at different salt concentrations. Biexponential fluorescence decays were observed at all salt concentrations, which were fitted to extract the relative proportion of the SRX state. A strong correlation was observed between the biochemical SRX state quantified by the SRX assay and structural information obtained from FRET efficiency calculated from the steady state fluorescence spectra (Figure 2B).

**Figure 2.**
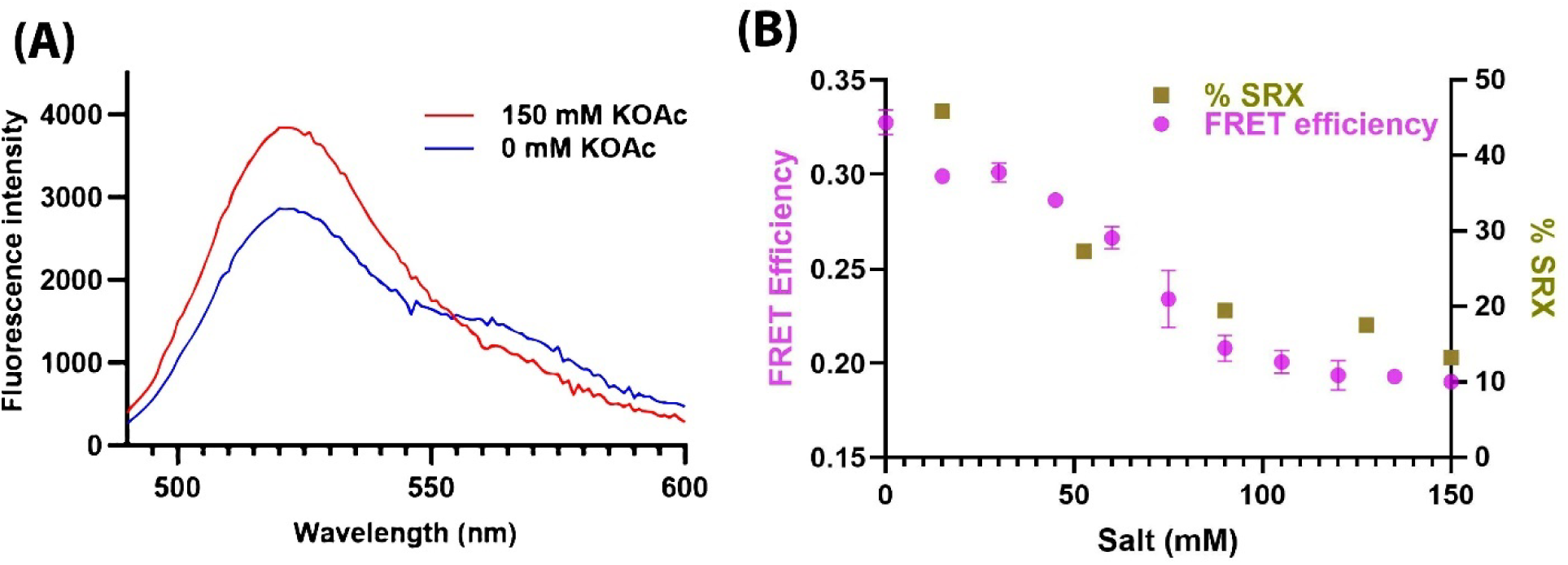
Steady state measurements with the RLC AF488-Cy3 ATP FRET sensor reports on structural changes in human β-cardiac 15-hep HMM due to changes in salt (KOAc) concentration. (A) Fluorescence emission spectra of 15-hep myosin with FRET sensor in the presence of 0 mM salt (blue curve) and 150 mM salt (red curve). (B) FRET efficiency calculated from the fluorescence emission spectra obtained at different salt concentrations is plotted along with the % SRX state obtained from the single-turnover assay.

### Proximal tail shortening results in disruption of the IHM state

Previous studies with smooth muscle myosin showed that at least 15 heptads of the proximal S2 are required to form the IHM OFF-state (Trybus et al., 1997). For β-cardiac myosin, a 15-hep HMM construct adopts the IHM conformation in solution (Grinzato et al., 2023; Rasicci et al., 2022), while an 8-hep HMM construct cannot, as judged by SRX and LSAR studies (Nandwani et al., 2024) (Figure 3A). Thus, 8-hep HMM behaves like short S1 (sS1, single headed myosin with only ELC bound) and 2-hep HMM constructs, neither of which is expected to form the IHM because the proximal S2 is either lacking (sS1) or not long enough for a stable head-tail interface to form (2-hep) (Anderson et al., 2018; Morck et al., 2022). Therefore, the 8-hep β-cardiac HMM should not show a salt-dependent change in FRET efficiency using our FRET sensor system.

**Figure 3.**
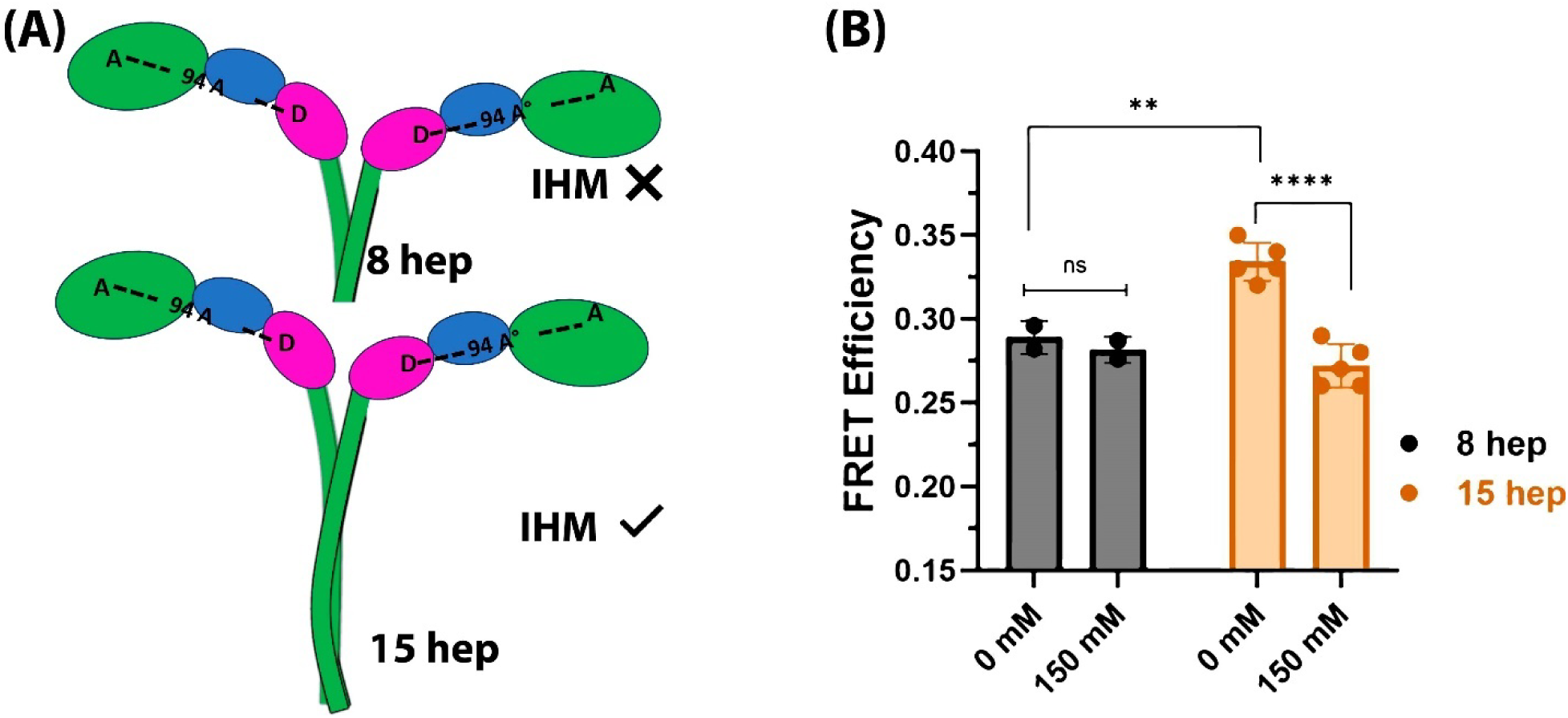
An 8-hep human β-cardiac HMM construct that cannot form the IHM state shows lower time-resolved FRET efficiency that is independent of salt concentration. (A) Schematic diagram showing difference between the 15-hep and 8-hep HMM constructs. The 15-hep HMM construct has been shown to form the folded back state while the 8-hep HMM construct cannot form the folded back state. (B) FRET efficiencies measured by time-resolved fluorescence for 8- and 15-hep β-cardiac HMM at 0 mM and 150 mM salt (KOAc) concentrations. The data for 15-hep HMM was collected from 3 different protein preparations measured across 4 different experiments and the data for 8-hep HMM was obtained from 2 different protein preparations across two different experiments.

We incorporated the donor and acceptor fluorophores in both 8-hep human β-cardiac HMM and 15-hep human β-cardiac HMM and measured the FRET efficiency of both by time-resolved fluorescence methodology. As shown in Figure 2B using steady-state FRET measurements of the 15-hep HMM construct, time-resolved FRET efficiency of the 15-hep HMM construct decreased when the salt concentration was increased to 150 mM (Figure 3B). For the 8-hep HMM construct, however, FRET efficiency remained unchanged when salt concentration was increased (Figure 3B). In the absence of salt, where the IHM state is most stabilized, the FRET efficiency for 15-hep HMM was significantly higher than 8-hep HMM. This result confirms that the 15-hep HMM construct can form the IHM configuration. The decrease in FRET efficiency for the 8-hep HMM construct indicates that the distance probed by the FRET sensor is higher in the 8-hep HMM construct, presumably due to the heads being open in this construct.

### The FRET sensor detects the effect of cardiomyopathy-causing mutations in β-cardiac myosin on the stability of the IHM state

Experiments from our lab and others (Craig & Padrón, 2022; Nag et al., 2017; Robert-Paganin et al., 2018; Spudich et al., 2024) have indicated that mutations in myosin that cause HCM result in destabilization of the IHM state, producing a pathological increase in force production due to a greater number of myosin heads interacting with actin. Most previous studies have evaluated the stability of the IHM state indirectly by quantitating the SRX state biochemically, assuming that the biochemical SRX state is a good proxy for the IHM state. However, as discussed above, this assumption does not always hold true. We used the FRET sensor to examine the effects of one HCM mutant (P710R) and one DCM mutant (E525K), in human β-cardiac myosin, on the levels of IHM, and we simultaneously measured the SRX levels. For these two mutants, the changes in SRX levels coincide with the FRET sensor results, as described below.

The P710R HCM mutation has been previously characterized extensively (Vander Roest et al., 2021),) and increased availability of myosin heads (measured by the LSAR assay and by decrease in SRX level) was the key hypercontractile change observed for this variant at the molecular level. The E525K DCM mutation has recently been shown to stabilize the SRX and IHM using biochemical assays (SRX and in vitro motility) and structural probes (TR-FRET and negative-staining EM) (Duno-Miranda et al., 2024; Rasicci et al., 2022), serving as a good control in our study.

We quantified the levels of SRX in wild type (WT) and both mutant variants of myosin, by performing the SRX assay as a function of salt concentration (Figures 4A). The P710R HCM mutation showed a reduced % SRX, whereas the E525K DCM mutation showed an increased % SRX at all salt concentrations tested, including 150 mM KOAc where less than 20% of molecules exist in the SRX state for WT and P710R mutant myosin (Figure 4A). Next, we measured FRET in the mutant and WT myosin 15-hep constructs with the FRET sensor, and the FRET efficiency was calculated by the change in D lifetime in the presence and absence of A using Equation 3. The FRET efficiency of the P710R mutant was lower than that of the WT myosin (Figure 4B, C) whereas the FRET efficiency of the E525K mutant was higher than the WT protein at both low (Figure 4B) and high (Figure 4C) salt concentrations. This observation indicates that the IHM state is populated more in the E525K DCM mutant and is populated less in the P710R HCM mutant, and that the IHM state of the E525K mutant is resistant to destabilization by 150 mM KOAc.

**Figure 4.**
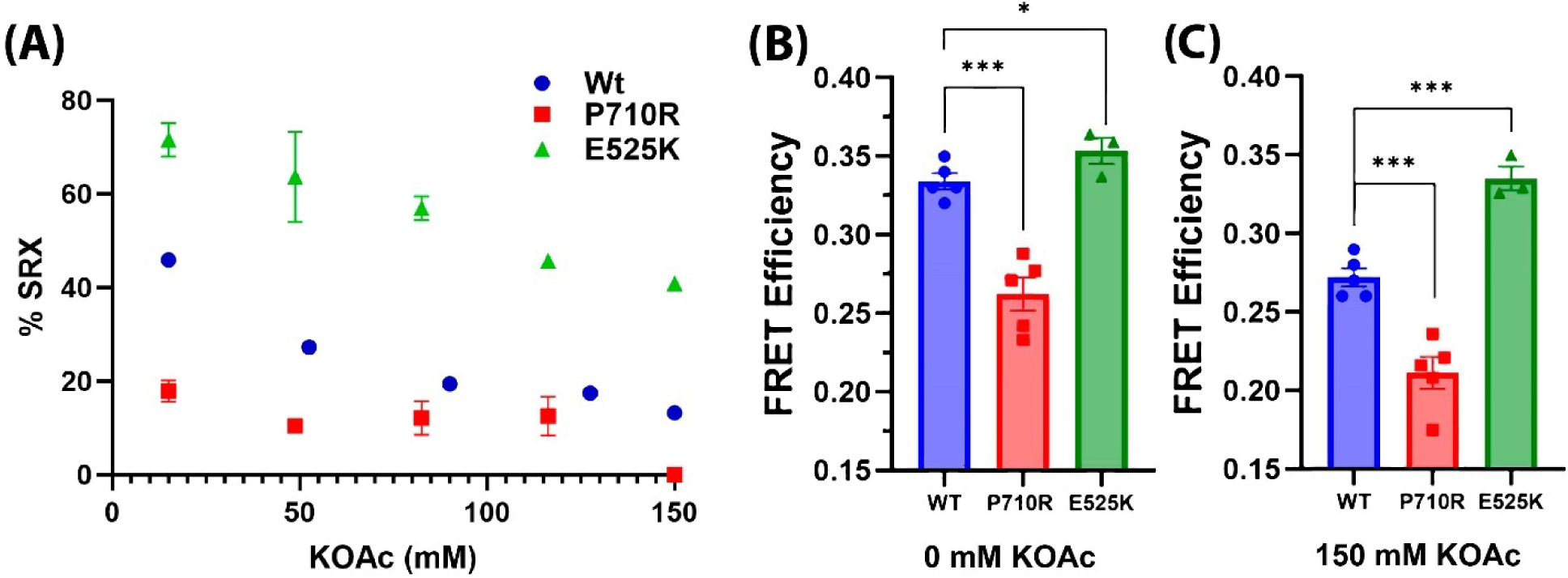
FRET sensor demonstrates altered SRX and FRET efficiencies due to cardiomyopathy-causing mutations in β-cardiac myosin. (A) Plot showing % SRX quantified by the SRX assay at different salt concentrations. Data shown is for 15-hep HMM constructs of WT human β-cardiac HMM (blue), HCM mutant P710R (red), and DCM mutant E525K (green). (B, C) Comparison of FRET efficiency measured by time-resolved fluorescence for WT, P710R and E525K 15hep β-cardiac HMM at 0 mM (B) and 150 mM (C) salt concentrations.

### The FRET sensor reliably reports on the IHM state in solution

The FRET sensor was tested by making several perturbations that are known to stabilize or destabilize the IHM state. In all cases, the theoretical calculated (expected) increase or decrease in FRET efficiency based on the near-atomic structure of the IHM state is in agreement with the experimentally determined FRET efficiency, as measured by the change in the lifetime of the donor fluorophore in the presence vs absence of the acceptor (Table 1). We compared these changes with the calculated FRET efficiency change for each experimental condition. To calculate the FRET efficiency, we used the results from the SRX assay and assumed that the observed change in the biochemically measured SRX fraction is due to a change in the structural IHM state. We used distances calculated from the atomic structure of the human β-cardiac IHM and the population of IHM indirectly obtained from SRX measurements to calculate expected FRET efficiency changes (Table 1, expected change in FRET efficiency).

**Table 1.**
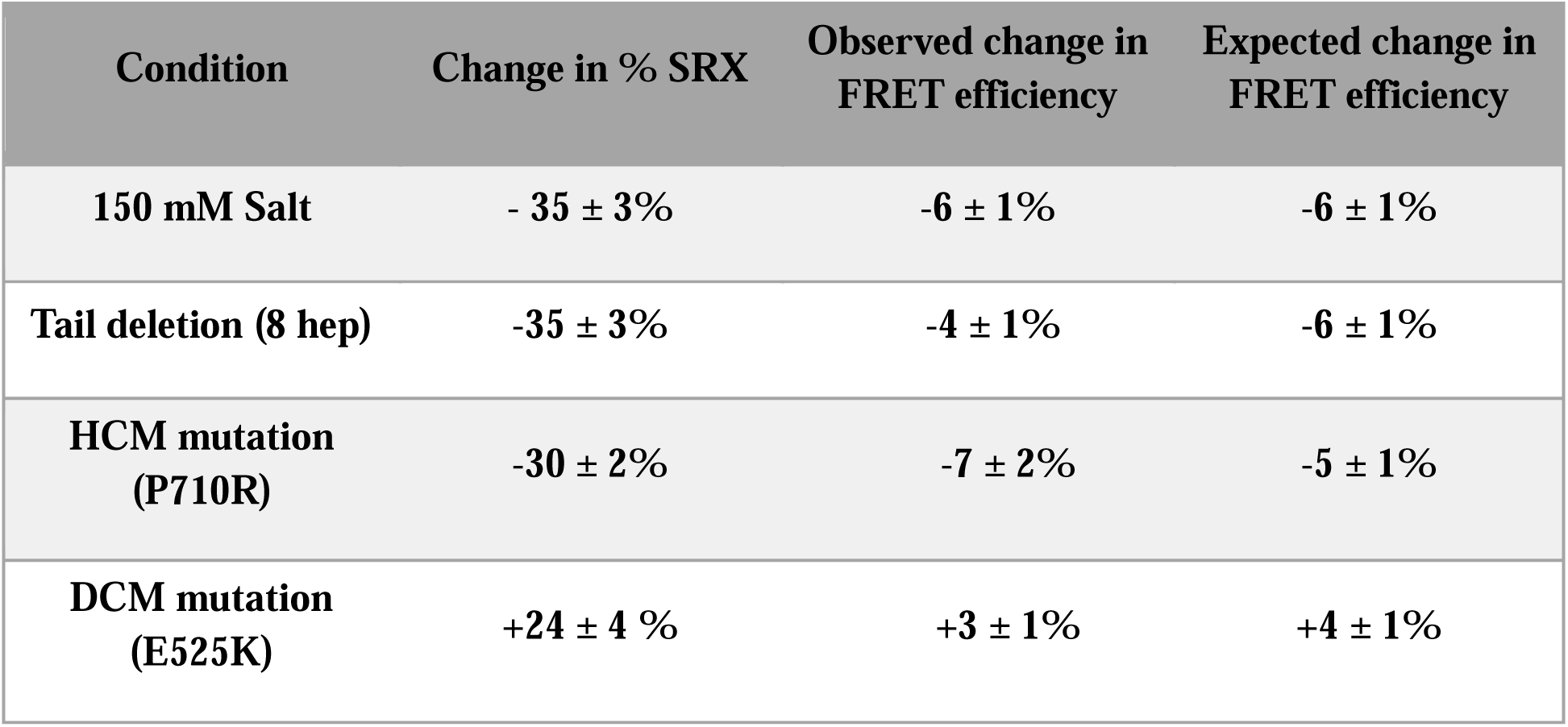
Comparison of experimentally observed changes in FRET efficiency (E) with theoretically calculated values. FRET efficiency and % SRX values are compared to those observed for WT 15-hep HMM at 0 mM salt. The FRET efficiency values were obtained from TR-FRET and %SRX values were obtained from SRX assay. Theoretical FRET efficiency was calculated from the distances observed in the near-atomic structure of the IHM state (PDB ID: 8ACT) using the R_0_ value of 67.6 Å and using the equation E = (1/1+(R_0_/R)^6^). The distances probed by the FRET sensor are indicated in Figure 1. For calculating the overall FRET efficiency in solution, it is assumed that % change SRX = % change IHM.

| Condition | Change in % SRX | Observed change in FRET efficiency | Expected change in FRET efficiency |
| --- | --- | --- | --- |
| 150 mM Salt | - 35 ± 3% | -6 ± 1% | -6 ± 1% |
| Tail deletion (8 hep) | -35 ± 3% | -4 ± 1% | -6 ± 1% |
| HCM mutation (P710R) | -30 ± 2% | -7 ± 2% | -5 ± 1% |
| DCM mutation (E525K) | +24 ± 4 % | +3 ± 1% | +4 ± 1% |

## Discussion

### Unique features of the present FRET sensor

FRET sensors for quantifying the IHM state were previously reported (Chu et al., 2021; Rasicci et al., 2022). The FRET sensor used here to detect the cardiac myosin IHM in solution is different in several respects from these two previously developed FRET sensors. The main difference is that our FRET sensor is the only one for which human β-cardiac myosin heavy chain was fully reconstituted with human ventricular ELC and RLC. The sensor reported by Chu *et al*. had donor and acceptor labels at position 105 of the two RLCs of bovine cardiac HMM. That sensor was able to detect FRET in the folded IHM state, but the distance in the open conformation was too long to show any FRET due to the short R_0_ of the FRET pair used. The second sensor, reported by Rasicci *et al*., measured FRET between GFP tags on the C terminus of the coiled coil tail of 15-hep myosin (donor) and Cy3-ATP (acceptor) bound in the nucleotide binding pocket of the motor head. Using that sensor, a strong correlation between the SRX and IHM state was observed as a function of ionic strength. Furthermore, the DCM-causing mutation E525K, which stabilizes the SRX state, showed an increase in FRET, consistent with an increase in IHM levels. However, the distances monitored by the Rasicci *et al*. FRET sensor (90-140 Å in the IHM and ≥250 Å in the open state) were larger than the R_0_ of the FRET pair used. As a result, FRET efficiency changes were very small, and, although the changes observed agreed qualitatively with the formation of IHM state, they could not be used to calculate distance changes.

In the case of our FRET sensor, the fluorophore was added to the C-terminus of the RLC for two reasons. First, the N-terminal region of the RLC is known to be involved in establishing the RLC-RLC interface that stabilizes the IHM (Grinzato et al., 2023), and phosphorylation of Ser 15 in the N-terminus is known to disrupt the IHM state *via* disruption of this interface (Dutta et al., 2023). AF488 carries a net negative charge, so we avoided modifying the N-terminus of RLC. Second, the distance between the C-terminus of the RLC and the nucleotide pocket in the free head and blocked head appears to be less than the distance from the N-terminus of the RLC. While the structures of neither the N-terminus nor C-terminus of the RLC are resolved in the high-resolution-cardiac 15-hep IHM structure (PDB ID: 8ACT), there is greater uncertainty about the N-terminus, since the first 19 amino acids of the N-terminus are missing, while the C-terminus is missing only the last 4 amino acids. The C-terminus of the RLC is probably closer to the motor head, making it likely that the distance is shorter between the nucleotide binding pocket and the C-terminus of the RLC. A shorter distance would allow greater energy transfer between D and A, resulting in higher FRET efficiency changes. The distances monitored here were in the optimal regime of the distances that can be quantified by FRET, which is between 0.5 and 1.5 R_0_ (Lakowicz, 1999; Stryer & Haugland, 1967). We were, therefore, able to compare the experimentally observed FRET changes with theoretically expected FRET changes.

### Confirmation that our FRET measurements are measuring levels of the structural IHM state

The effects of HCM mutations on the stability of the IHM has been inferred from a range of assays of which the indirect biochemical SRX assay is the most frequently used. Unlike the SRX assay, the FRET approach reported here more directly probes the IHM conformation of cardiac myosin in solution. This FRET approach robustly monitors the dynamics of IHM, indicated by its sensitivity to both solution ionic strength and requirement of a long enough proximal S2 tail length to allow stabilization of the IHM. Sensitivity to ionic strength was reported for only one of the previously designed FRET sensors (Rasicci et al., 2022), while the tail-length dependence was not tested for either of them (Chu et al., 2021; Rasicci et al., 2022).

Our FRET sensor was further tested for reporting on the population of the IHM state using two pathogenic mutations in myosin that have been extensively characterized. The P710R HCM mutant variant has previously been shown to destabilize the biochemical SRX state as well as the functional autoinhibited state inferred from the LSAR assay (Vander Roest et al., 2021). In fact, increased availability of myosin heads is the primary change at the molecular level contributing to the hypercontractile disease phenotype of this mutation, a phenotype that has been validated in P710R-expressing induced pluripotent stem cell-derived cardiomyocytes (Vander Roest et al., 2021). Our FRET sensor shows directly that the destabilization of SRX observed in the absence of actin is indeed well correlated with the destabilization of the IHM (Table 1). For the DCM-causing E525K mutation, an increase in the SRX fraction correlated well with the increase in FRET efficiency at both low and high salt, similar to the qualitative findings using the head-tail FRET sensor reported earlier by Rasicci *et al*. (Rasicci et al., 2022). E525K is the only SRX- and IHM-stabilizing mutation reported thus far, and while it serves as an excellent control here, it also shows that our sensor is very versatile, with a large enough dynamic range to reliably measure both destabilization and stabilization of the IHM.

### Factors to consider when using the FRET sensor described here

The FRET changes observed for different perturbations of the IHM ⇔ open state equilibrium were very small, <10%. We were able to robustly measure such small changes by using highly homogeneous myosin that contained 100% fluorescently labeled RLC (Supplementary Figure S1). It was also necessary to use a high-resolution technique such as time-resolved FRET to accurately calculate the changes in FRET efficiency at different conditions. We were also able to observe FRET efficiency changes using steady-state fluorescence measurements, although the values of FRET efficiency were slightly different between the two methods. This is not surprising, since the steady-state fluorescence intensity of Cy3 can be erroneous due to slight overlap in the emission spectra of AF488 and Cy3. We also observed a significant direct excitation of the acceptor Cy3-ATP at 470 nm wavelength used for donor excitation. This direct excitation is corrected by subtracting a corresponding blank solution containing only Cy3-ATP without the AF488 labeled protein. Because of these issues with steady-state FRET measurements, the calculations for Table 1 were done using time-resolved fluorescence data. The FRET efficiencies calculated from time-resolved lifetime waveforms are very accurate since they are calculated by measuring the lifetime of the donor in the presence and absence of acceptor, independent of environmental factors. The use of time-corelated single photon counting (TCSPC) methodology to determine fluorescence lifetime wavefronts of the AF488 label at a very high signal to noise enabled reliable detection of even a 1% change in the lifetime(Lakowicz, 1983).

The Cy3-ATP acceptor used in the FRET sensor is not covalently attached to the myosin and it undergoes cycling as myosin consumes ATP, resulting in release of the label. Myosin also undergoes conformational changes as it cycles Cy3-ATP, necessitating the need for fast-mixing devices to make such FRET measurements, as was the case with the Rasicci *et al*. sensor (Rasicci et al., 2022). To allow for an equilibrium FRET analysis, a 100-fold molar excess of Cy3-ATP was used to ensure that the myosin catalytic heads were always bound to the nucleotide. Since the of inorganic phosphate (P_i_) is the rate-limiting step in the chemomechanical cycle of myosin, in the presence of excess ATP most of the myosin heads are in the pre-powerstroke state (PPS), bound with ADP and P_i_, and the distance between the AF488-Cy3-ATP FRET pair corresponds to the FH distance in both heads in most molecules. In the IHM state, the near-atomic resolution structure of the IHM clearly shows ADP and P_i_ bound at the nucleotide binding pocket, and the two motor domains are in a classical PPS conformation. The kink in the pliant region of the BH brings the AF488-Cy3 FRET pair closer together, resulting in a change in FRET efficiency as solution conditions are varied. The FRET measurements performed in the presence of excess ATP therefore reflects the equilibrium between the PPS state of myosin and the IHM state.

We used a reported R_0_ value of 67.6 Å for the D-A pair (Muretta et al., 2015) used in the current study to calculate the expected FRET changes from the distances measured from the atomic structure of the IHM. The reported R_0_ value assumes that the fluorophores are freely rotating (hence an assumed orientation factor κ^2^ of 2/3) and that the overlap integral (the region of overlap between the donor emission spectrum and the acceptor absorption spectrum for the free fluorophore) remains unchanged. The previous study has addressed some of these issues, concluding that the AF488-Cy3 FRET pair is very sensitive to the distance changes between the myosin lever arm and nucleotide binding pocket (Muretta et al., 2015), and this FRET pair was later successfully used to monitor the structural effects of myosin modulators omecamtiv mecarbil (Rohde et al., 2017) and mavacamten (Rohde et al., 2018) on bovine cardiac myosin.

The FRET sensor described here has a D-A pair on each head of the 15-hep myosin construct. Each donor can undergo FRET with the acceptor bound on the same head or on the other head. Of the four possible combinations of distances, only two are indicated in the schematic in Figure 1A, the distance between the BH Cy3-ADP and BH RLC, and a longer distance between the FH Cy3-ADP and FH RLC. The two other distances not indicated in Figure 1A are the distance between the BH RLC and FH Cy3-ADP and the distance between the FH RLC and BH Cy3-ADP. These two distances are very similar to the two distances indicated in Figure 1A, and although there are 4 possible D-A pairs, distance-wise there are only two possible distances.

### Caution must be used in interpretations from SRX measurements

The IHM state has been assumed to be the structural basis of the biochemical SRX state in muscle fibers and in purified myosin preparations (Craig & Padrón, 2022). Several studies show that caution is warranted when using the SRX assay as a sole measure of the IHM state (Chu et al., 2021; Jani et al., 2024; Ma et al., 2023; Morck et al., 2022). The structural origins of biochemical SRX state are not yet completely understood and it is not trivial to predict whether a particular mutation in β-cardiac myosin will stabilize/destabilize the IHM state (Nandwani et al., 2024). Thus, caution must be used when interpreting measurements using only the SRX assay for levels of SRX or for equating changes in IHM levels to changes in low angle X-ray scattering or polarization of fluorescence measurements in fibers. In the current study we observe an excellent agreement between the SRX state and IHM state when we consider the overall FRET efficiency changes (Table 1). Given all these issues, it is highly recommended to validate the results of SRX assay with other biochemical methods such as LSAR and FRET assays.

In the FRET assay in this study, the decays are interpreted in terms of resolved distance distributions, which appear to correlate well with known SRX changes. The various controls tested, such as change in ionic strength and variation in proximal S2 length, conclusively showed that the distance between the C-terminus of RLC and the nucleotide binding pocket is a very good structural indicator of the IHM state. In contrast, fiber-based structural methods such as X-ray diffraction and fluorescence polarization are not direct indicators of the IHM state. Movement of the proximal S2 tail along with one or both myosin heads in the IHM state away from the filament can lead to a disruption of the quasihelical arrangement of myosin heads, probed by X-ray diffraction, and to disorder that can result in depolarization of fluorescence. Several fundamental issues regarding the dynamics of the IHM state still need to be explored. For example, fluorescence polarization and X-ray diffraction studies show that the IHM state undergoes dynamics at the timescale of the chemo-mechanical cycle of myosin(Irving, 2017, 2024). How can such a dynamic structural state give rise to a slow SRX state is unclear. Future FRET studies aimed at understanding the dynamics of the IHM state will help to resolve these issues.

## AUTHOR CONTRIBUTIONS

RRG conceptualized and designed the study, performed protein expression, purification and labeling, steady state and time-resolved FRET and single turnover experiments, and analyzed the data. PG performed time-resolved FRET experiments and analyzed the data. NN performed protein purification and initial SRX measurements of E525K mutant. AD and SY performed adenoviral production. RRG wrote the original draft, and PG, NN, AD, SY, OR, DDT, JAS, and KMR contributed to editing and reviewing of the manuscript. DDT, JAS and KMR supervised the study and acquired funding.

## ACKNOWLEDGEMENTS

The authors thank all members of the Spudich/Ruppel lab for discussions, training and support, and Samantha Yuen of the Thomas lab for help with the time-resolved fluorescence measurements. We thank Dr. Anne Houdusse for helping with the distance calculations. This work was funded by NIH grants HL117138 and 2GM033289 to J.A.S. and K.M.R. and postdoctoral fellowships from American Heart Association to R.R.G. (Award ID 23POST1027175) and N.N. (Award ID 22POST908934). All the time-resolved fluorescence measurements were performed at the Biophysical Technology Centre (BTC), directed by Dr. Robyn Rebbeck at the University of Minnesota.

## Supplementary Information (SI)

### Distances probed by the FRET sensor

In our FRET sensor the Donor dye is attached to a cysteine incorporated at the C-terminus of RLC (Cys-RLC). The last four residues after Glycine 162 of the C-terminus of the RLC were not resolved in the structure. The distances indicated in Figure 1A and 1B are measured from Gly 162 to the OH group of the ADP bound in the nucleotide binding pocket to which acceptor Cy3 was attached. The four residues in the C-terminus followed by a Cysteine will add an additional ∼ 15 Å to the measured distance. In addition to this uncertainty both Donor and acceptor fluorophores have linkers attached to them. The donor AF488 dye has a C5 linker that adds additional ∼15 Å to the distance and Acceptor Cy3 attached to the OH group of ribose through a linker of consisting of 12 covalent bonds which adds a distance of ∼ 30 Å. Assuming that the linkers can sample different conformations radially around the site of attachment (green and red circles in Figure 1C) the possible distances for D-A pair were shown in Figure 1C. These linkers add an additional 45 Å to the actual distance distributions centered around 69 Å for BH and 94 Å for the FH. Not all orientations of the fluorophore are equally probable due to steric clashes with the different structural elements in the protein and only few of the rotamers of the dye linker pair are energetically favorable. The actual DA distance distributions were difficult to measure experimentally. Given the length of these linkers and the rough estimate of distances from the atomic structure It is likely that the DA distance can be as short as ∼20 Å for BH and ∼49 Å for FH and as long as ∼114 Å for BH and ∼ 139 Å for FH. The distance distributions can result in different rates of energy transfer between DA pairs resulting in different time scales of decay of the donor fluorescence in the presence of acceptor which is measured experimentally.

**Figure S1.**
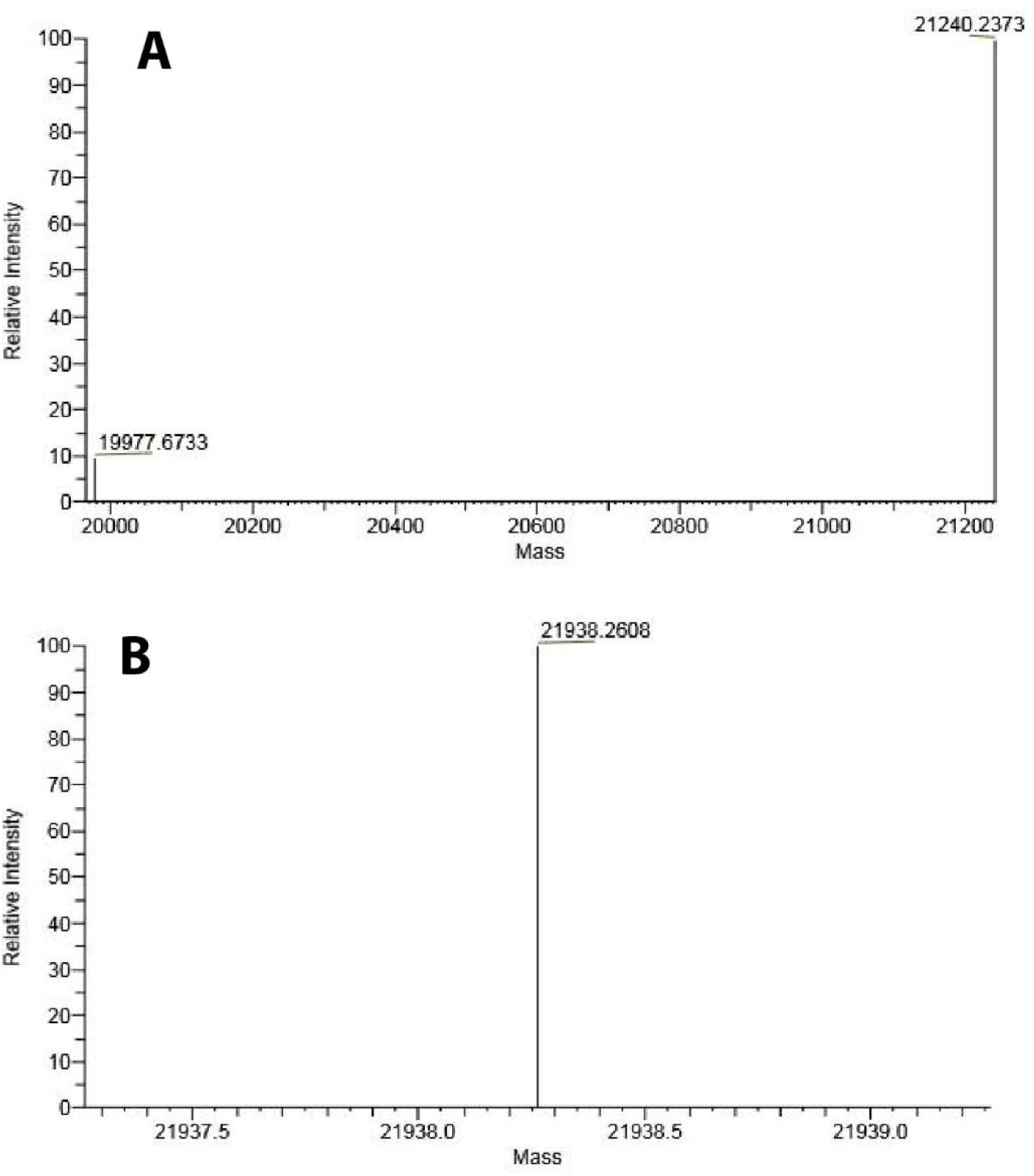
Assessing the extent of labeling of the Cys-RLC with AF488 by mass spectrometry. (A) Mass spectrum showing mass of unlabeled human Cys-RLC. (B) Mass spectrum showing mass of Cys-RLC with AF488 label attached. The increase in mass of 698 Da corresponds to attachment of one AF488 label to the protein.

**Figure S2.**
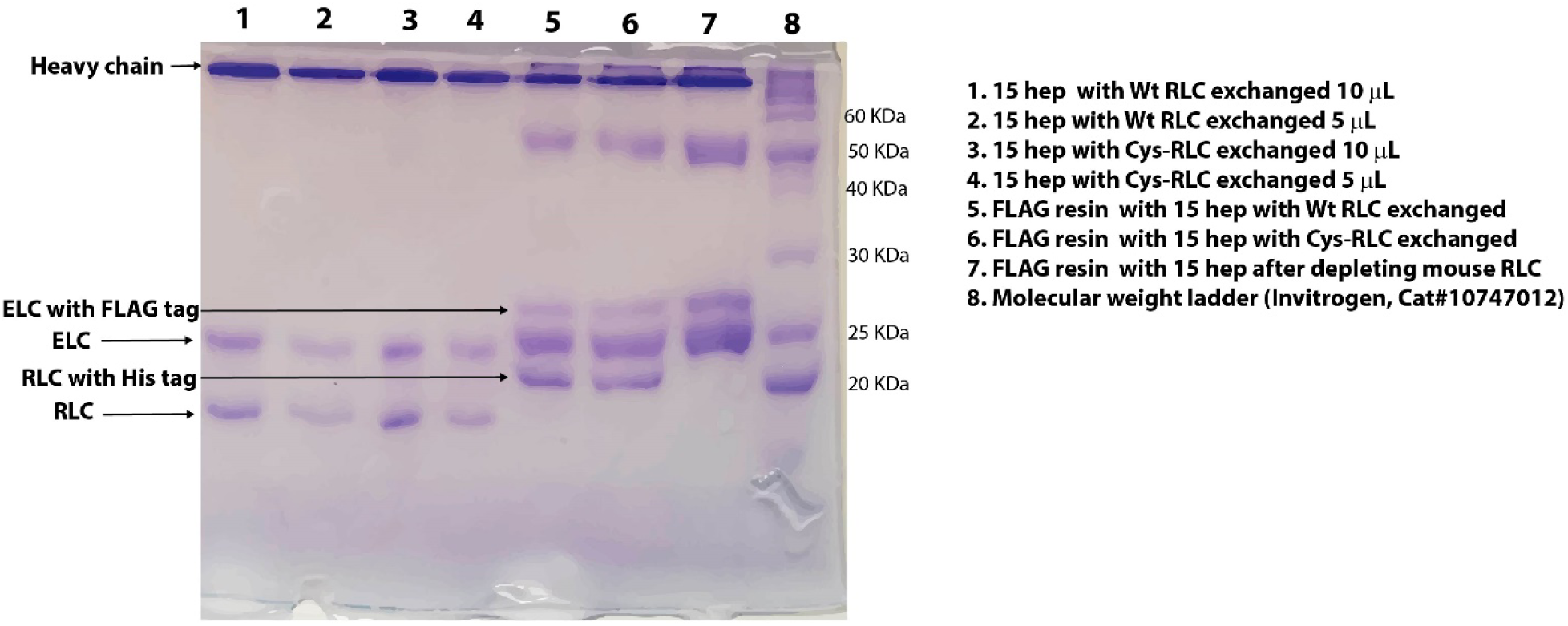
**Purity and extent of RLC exchange of 15-hep HMM assessed by 15% SDS PAGE. Clarified lysate from** C2C12 cells expressing 15-hep HMM heavy chain and FLAG-tagged human ELC was subjected to affinity chromatography using anti-FLAG resin followed by ion exchange chromatography using a Q Sepharose column. FLAG resin after depleting the endogenous mouse RLC was loaded in lane 7. The resin was split into two parts and one part exchanged with WT RLC and the other part exchanged with Cys-RLC (see Materials and Methods), and the samples were loaded in lanes 5 and 6 respectively. The bands at molecular weight 50 and 25 KDa in lanes 5-7 correspond to heavy chain and light chain of anti-FLAG antibody of the resin. The protein was eluted off the FLAG resin by treating with a TEV protease and further purified by Q Sepharose chromatography which gets rid of the full-length endogenous mouse skeletal myosin heavy chain (band above 15 hep HMM heavy chain in lanes 5-7) as well as the TEV protease. The purified proteins were loaded in lanes 1-4 showing bands corresponding to the WT 15-hep HMM heavy chain and the human ventricular ELC and Cys-RLC.

**Figure S3.**
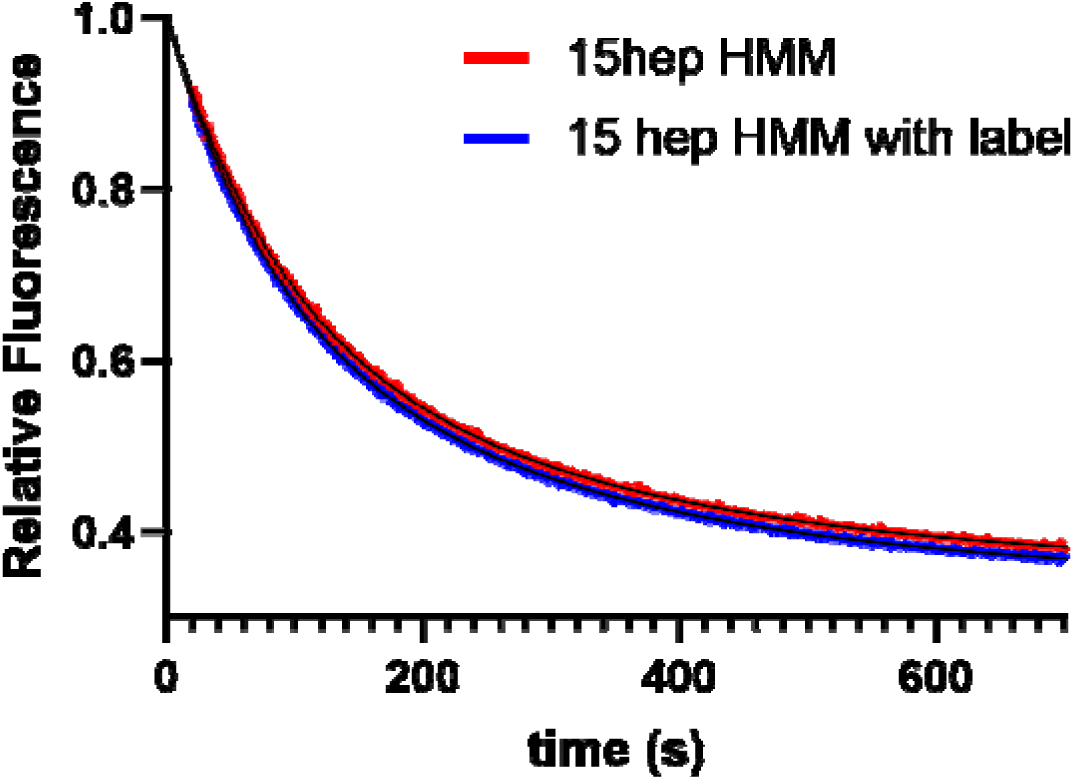
Labeling of the Cys-RLC does not alter single nucleotide turnover kinetics of WT β-cardiac 15-hep HMM. Kinetic traces are shown for WT 15-hep HMM (red) and WT 15-hep HMM with Cys-RLC labeled with AF488 (blue) at 15 mM salt. Both kinetic traces were fit to a double exponential equation, and the fits to the data are shown in black lines. The fit yielded rate constants of 0.01 s^-1^ (54% amplitude, DRX) and 0.003 s^-1^ (46% amplitude, SRX) for the unlabeled protein. The fit to the labeled protein trace yielded the same rate constants of 0.01 s^-1^ (50% amplitude, DRX) and 0.003 s^-1^ (50% amplitude, SRX).

**Figure S4.**
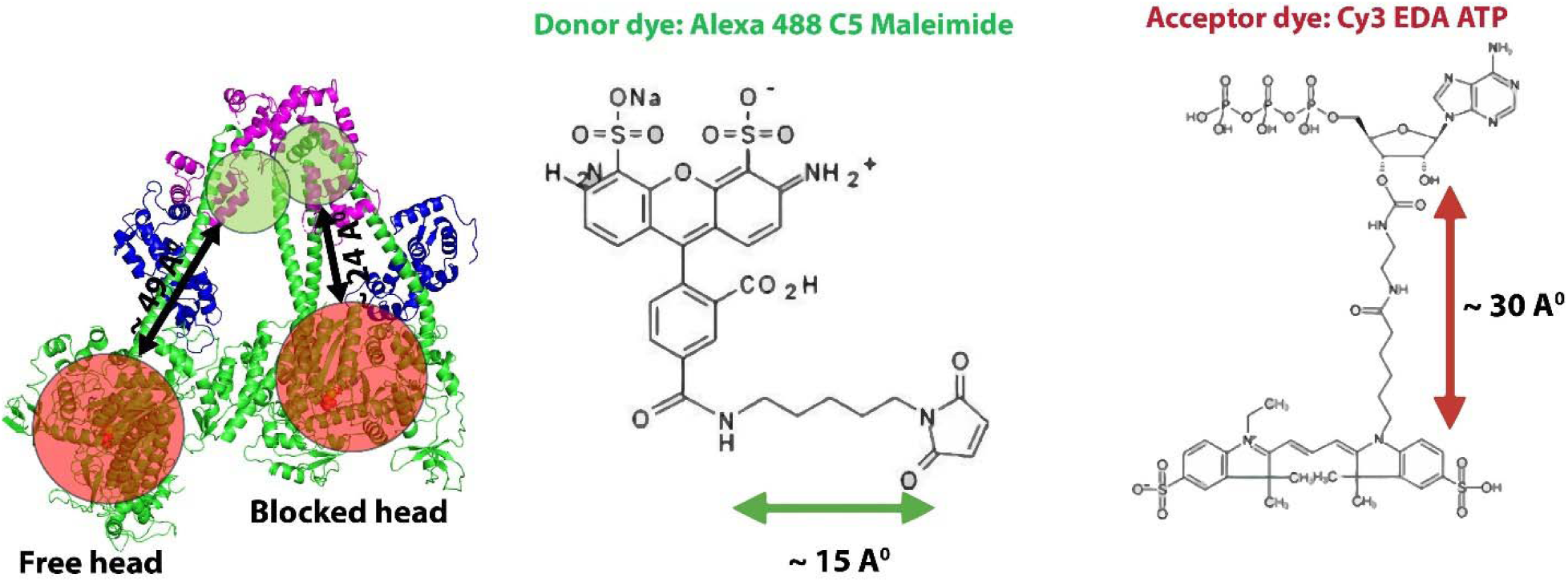
Estimation of distances probed by the FRET sensor. Schematic diagram of the IHM state of human β-cardiac 15-hep HMM showing the distances sampled by donor and acceptor fluorophores (left panel). Both the dyes used in the current FRET sensor have linkers to keep the probe away from the protein and to allow free rotation of fluorophore. The C5 linker on AF488 adds a distance of 15 Å (shown as green circle on the folded back state structure) the EDA linker on Cy3 ATP adds a distance of ∼ 30 Å. Assuming the linkers sample all the available conformations, the distance between the blocked head RLC and its nucleotide binding pocket can approach as close as ∼ 24 Å, while the distance for the free head can approach ∼ 49 Å.

**Table S1.**
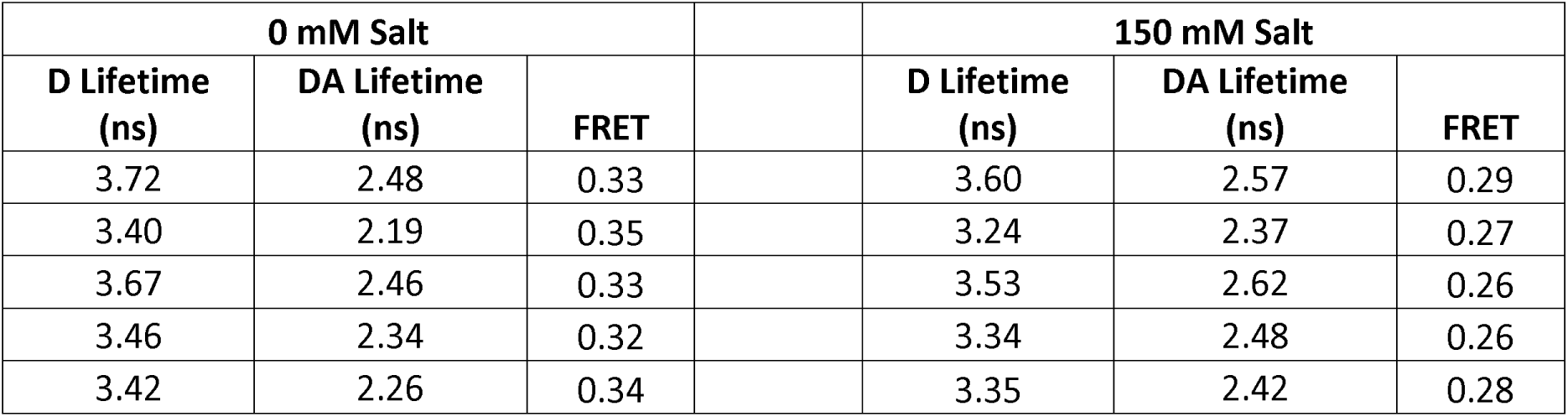
Average fluorescence lifetime values of WT 15-hep β-cardiac HMM. Amplitude weighted average lifetime values obtained at 0 and 150 mM salt concentrations. These values represent three independent protein preparations measured across 5 different experiments. FRET efficiency values shown in the table were used to generate Figure 3B.

**Table S2.**
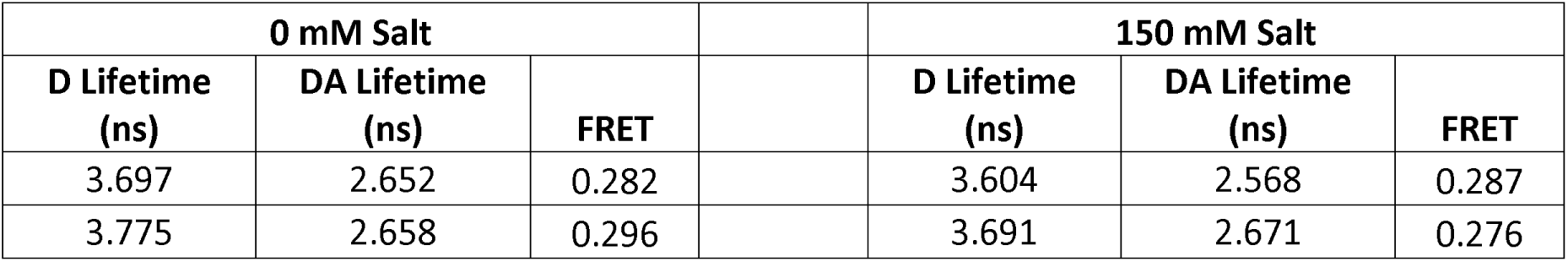
Average fluorescence lifetime values of WT 8 hep β-cardiac HMM. Amplitude weighted average lifetime values obtained at 0 mM and 150 mM salt concentrations. These values represent two independent protein preparations measured across two different experiments. FRET efficiency values shown in above table were used to generate Figure 3B.

| 0 mM Salt |  |  |  | 150 mM Salt |  |  |
| --- | --- | --- | --- | --- | --- | --- |
| D Lifetime<br>(ns) | DA Lifetime<br>(ns) | FRET |  | D Lifetime<br>(ns) | DA Lifetime<br>(ns) | FRET |
| 3.697 | 2.652 | 0.282 |  | 3.604 | 2.568 | 0.287 |
| 3.775 | 2.658 | 0.296 |  | 3.691 | 2.671 | 0.276 |

**Table S3.**
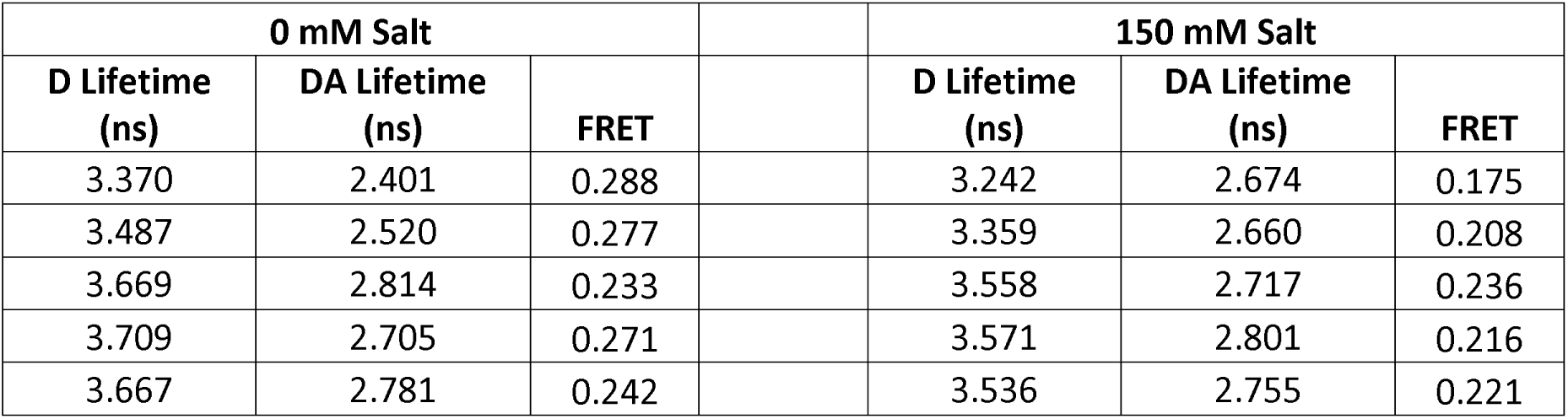
Fluorescence lifetime values of 15-hep β-cardiac HMM with the HCM mutation P710R. Amplitude weighted average lifetime values obtained at 0 mM and 150 mM salt concentrations. These values represent 2 independent protein preparations measured across 3 different experiments. In two of the experiments two additional technical replicates were obtained. For technical replicates the samples were prepared by diluting the stock protein into an appropriate buffer. FRET efficiencies values from this table were used to generate Figure 4B and 4C.

**Table S4.**
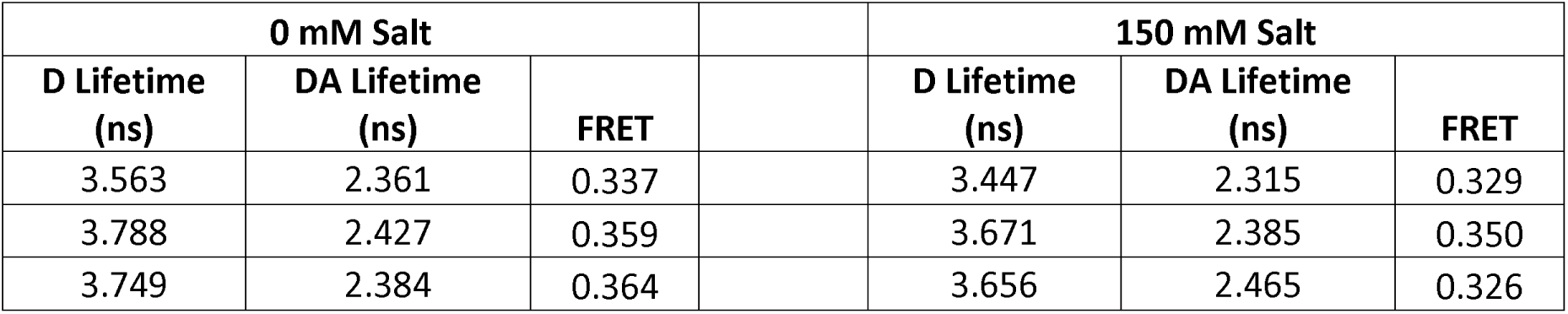
Fluorescence lifetime values of 15 hep β-cardiac HMM with the DCM mutation E525K. Amplitude weighted average lifetime values obtained at 0 mM and 150 mM salt concentrations. These values represent 2 independent protein preparations measured across 3 different experiments. FRET efficiencies values from this table were used to generate Figure 4B and 4C.

| 0 mM Salt |  |  |  | 150 mM Salt |  |  |
| --- | --- | --- | --- | --- | --- | --- |
| D Lifetime (ns) | DA Lifetime (ns) | FRET |  | D Lifetime (ns) | DA Lifetime (ns) | FRET |
| 3.563 | 2.361 | 0.337 |  | 3.447 | 2.315 | 0.329 |
| 3.788 | 2.427 | 0.359 |  | 3.671 | 2.385 | 0.350 |
| 3.749 | 2.384 | 0.364 |  | 3.656 | 2.465 | 0.326 |

**Table S5.**
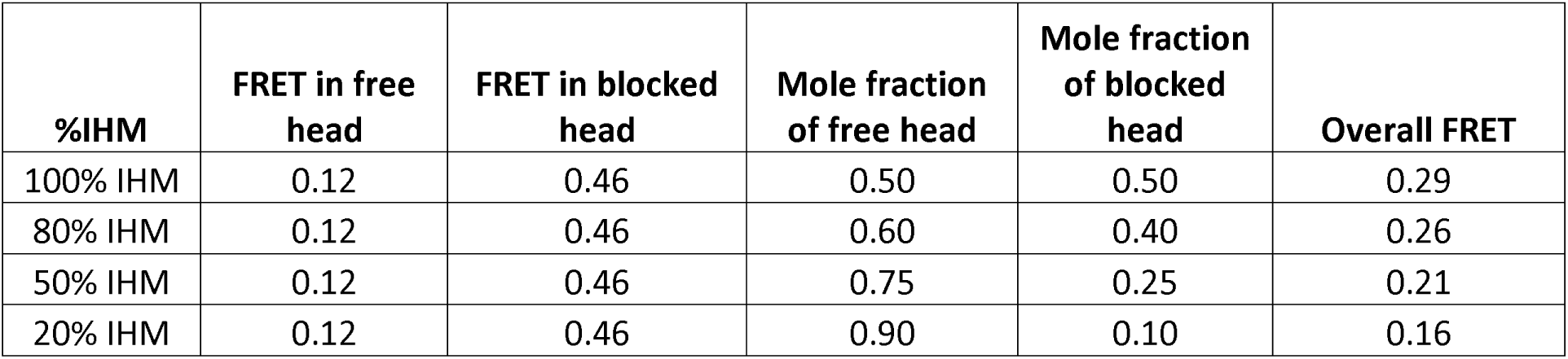
Theoretical FRET efficiencies calculated for different proportions of IHM present in solution. The distance between the C-terminus of the RLC and the hydroxyl group on the ribose moiety of the ADP is derived from the IHM structure (PDB ID: 8ACT, see Fig 1B). The distances for the free head and blocked head were used to calculate the FRET efficiency using an R_0_ value of 67.6 Å.

