## Supplementary Information for "A FRET assay to quantitate levels of the human β-cardiac myosin interacting heads motif based on its near-atomic resolution structure"

**Supplementary Information (SI)**

**Distances probed by the FRET sensor:**

In our FRET sensor the Donor dye is attached to a cysteine incorporated at the C-terminus of RLC (Cys-RLC). The last four residues after Glycine 162 of the C-terminus of the RLC were not resolved in the structure. The distances indicated in Figure 1A and 1B are measured from Gly 162 to the OH group of the ADP bound in the nucleotide binding pocket to which acceptor Cy3 was attached. The four residues in the C-terminus followed by a Cysteine will add an additional ~ 15 Å to the measured distance. In addition to this uncertainty both Donor and acceptor fluorophores have linkers attached to them. The donor AF488 dye has a C5 linker that adds additional ~15 Å to the distance and Acceptor Cy3 attached to the OH group of ribose through a linker of consisting of 12 covalent bonds which adds a distance of ~ 30 Å. Assuming that the linkers can sample different conformations radially around the site of attachment (green and red circles in Figure 1C) the possible distances for D-A pair were shown in Figure 1C. These linkers add an additional 45 Å to the actual distance distributions centered around 69 Å for BH and 94 Å for the FH. Not all orientations of the fluorophore are equally probable due to steric clashes with the different structural elements in the protein and only few of the rotamers of the dye linker pair are energetically favorable. The actual DA distance distributions were difficult to measure experimentally. Given the length of these linkers and the rough estimate of distances from the atomic structure It is likely that the DA distance can be as short as ~20 Å for BH and ~49 Å for FH and as long as ~114 Å for BH and ~ 139 Å for FH. The distance distributions can result in different rates of energy transfer between DA pairs resulting in different time scales of decay of the donor fluorescence in the presence of acceptor which is measured experimentally.

| **0 mM Salt** | | |  | **150 mM Salt** | | |
| --- | --- | --- | --- | --- | --- | --- |
| **D Lifetime (ns)** | **DA Lifetime (ns)** | **FRET** |  | **D Lifetime (ns)** | **DA Lifetime (ns)** | **FRET** |
| 3.72 | 2.48 | 0.33 |  | 3.60 | 2.57 | 0.29 |
| 3.40 | 2.19 | 0.35 |  | 3.24 | 2.37 | 0.27 |
| 3.67 | 2.46 | 0.33 |  | 3.53 | 2.62 | 0.26 |
| 3.46 | 2.34 | 0.32 |  | 3.34 | 2.48 | 0.26 |
| 3.42 | 2.26 | 0.34 |  | 3.35 | 2.42 | 0.28 |

**Table S1. Average fluorescence lifetime values of WT 15-hep β-cardiac HMM.** Amplitude weighted average lifetime values obtained at 0 and 150 mM salt concentrations. These values represent three independent protein preparations measured across 5 different experiments. FRET efficiency values shown in the table were used to generate Figure 3B.

| **0 mM Salt** | | |  | **150 mM Salt** | | |
| --- | --- | --- | --- | --- | --- | --- |
| **D Lifetime (ns)** | **DA Lifetime (ns)** | **FRET** |  | **D Lifetime (ns)** | **DA Lifetime (ns)** | **FRET** |
| 3.697 | 2.652 | 0.282 |  | 3.604 | 2.568 | 0.287 |
| 3.775 | 2.658 | 0.296 |  | 3.691 | 2.671 | 0.276 |

**Table S2. Average fluorescence lifetime values of WT 8 hep β-cardiac HMM.** Amplitude weighted average lifetime values obtained at 0 mM and 150 mM salt concentrations. These values represent two independent protein preparations measured across two different experiments. FRET efficiency values shown in above table were used to generate Figure 3B.

| **0 mM Salt** | | |  | **150 mM Salt** | | |
| --- | --- | --- | --- | --- | --- | --- |
| **D Lifetime (ns)** | **DA Lifetime (ns)** | **FRET** |  | **D Lifetime (ns)** | **DA Lifetime (ns)** | **FRET** |
| 3.370 | 2.401 | 0.288 |  | 3.242 | 2.674 | 0.175 |
| 3.487 | 2.520 | 0.277 |  | 3.359 | 2.660 | 0.208 |
| 3.669 | 2.814 | 0.233 |  | 3.558 | 2.717 | 0.236 |
| 3.709 | 2.705 | 0.271 |  | 3.571 | 2.801 | 0.216 |
| 3.667 | 2.781 | 0.242 |  | 3.536 | 2.755 | 0.221 |

**Table S3. Fluorescence lifetime values of 15-hep β-cardiac HMM with the HCM mutation P710R.** Amplitude weighted average lifetime values obtained at 0 mM and 150 mM salt concentrations. These values represent 2 independent protein preparations measured across 3 different experiments. In two of the experiments two additional technical replicates were obtained. For technical replicates the samples were prepared by diluting the stock protein into an appropriate buffer. FRET efficiencies values from this table were used to generate Figure 4B and 4C.

| **0 mM Salt** | | |  | **150 mM Salt** | | |
| --- | --- | --- | --- | --- | --- | --- |
| **D Lifetime (ns)** | **DA Lifetime (ns)** | **FRET** |  | **D Lifetime (ns)** | **DA Lifetime (ns)** | **FRET** |
| 3.563 | 2.361 | 0.337 |  | 3.447 | 2.315 | 0.329 |
| 3.788 | 2.427 | 0.359 |  | 3.671 | 2.385 | 0.350 |
| 3.749 | 2.384 | 0.364 |  | 3.656 | 2.465 | 0.326 |

**Table S4. Fluorescence lifetime values of 15 hep β-cardiac HMM with the DCM mutation E525K.** Amplitude weighted average lifetime values obtained at 0 mM and 150 mM salt concentrations. These values represent 2 independent protein preparations measured across 3 different experiments. FRET efficiencies values from this table were used to generate Figure 4B and 4C.

| **%IHM** | **FRET in free head** | **FRET in blocked head** | **Mole fraction of free head** | **Mole fraction of blocked head** | **Overall FRET** |
| --- | --- | --- | --- | --- | --- |
| 100% IHM | 0.12 | 0.46 | 0.50 | 0.50 | 0.29 |
| 80% IHM | 0.12 | 0.46 | 0.60 | 0.40 | 0.26 |
| 50% IHM | 0.12 | 0.46 | 0.75 | 0.25 | 0.21 |
| 20% IHM | 0.12 | 0.46 | 0.90 | 0.10 | 0.16 |

**Table S5. Theoretical FRET efficiencies calculated for different proportions of IHM present in solution.** The distance between the C-terminus of the RLC and the hydroxyl group on the ribose moiety of the ADP is derived from the IHM structure (PDB ID: 8ACT, see Fig 1B). The distances for the free head and blocked head were used to calculate the FRET efficiency using an R_0_ value of 67.6 Å.

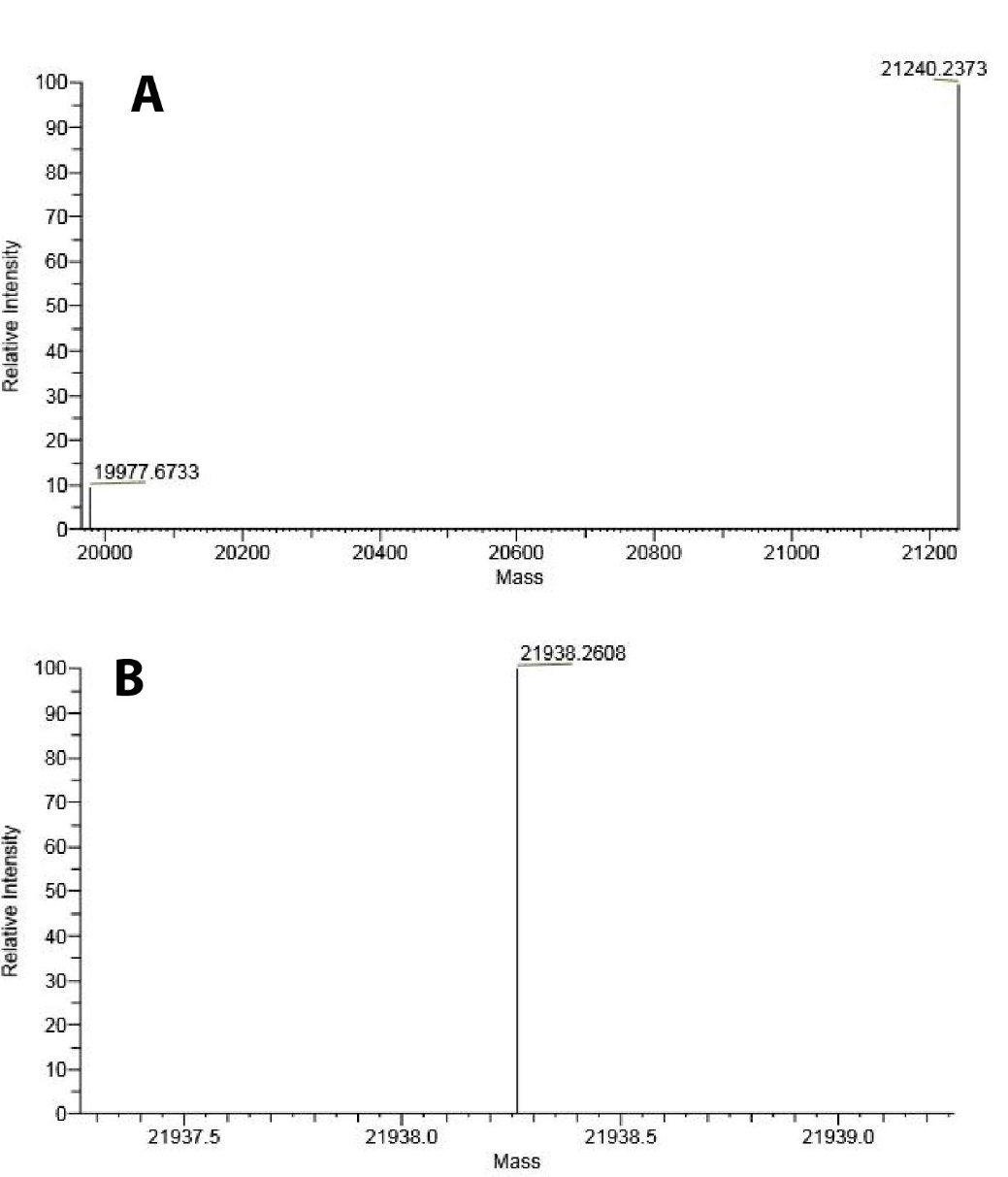

**Figure S1. Assessing the extent of labeling of the Cys-RLC with AF488 by mass spectrometry.** (A) Mass spectrum showing mass of unlabeled human Cys-RLC. (B) Mass spectrum showing mass of Cys-RLC with AF488 label attached. The increase in mass of 698 Da corresponds to attachment of one AF488 label to the protein.

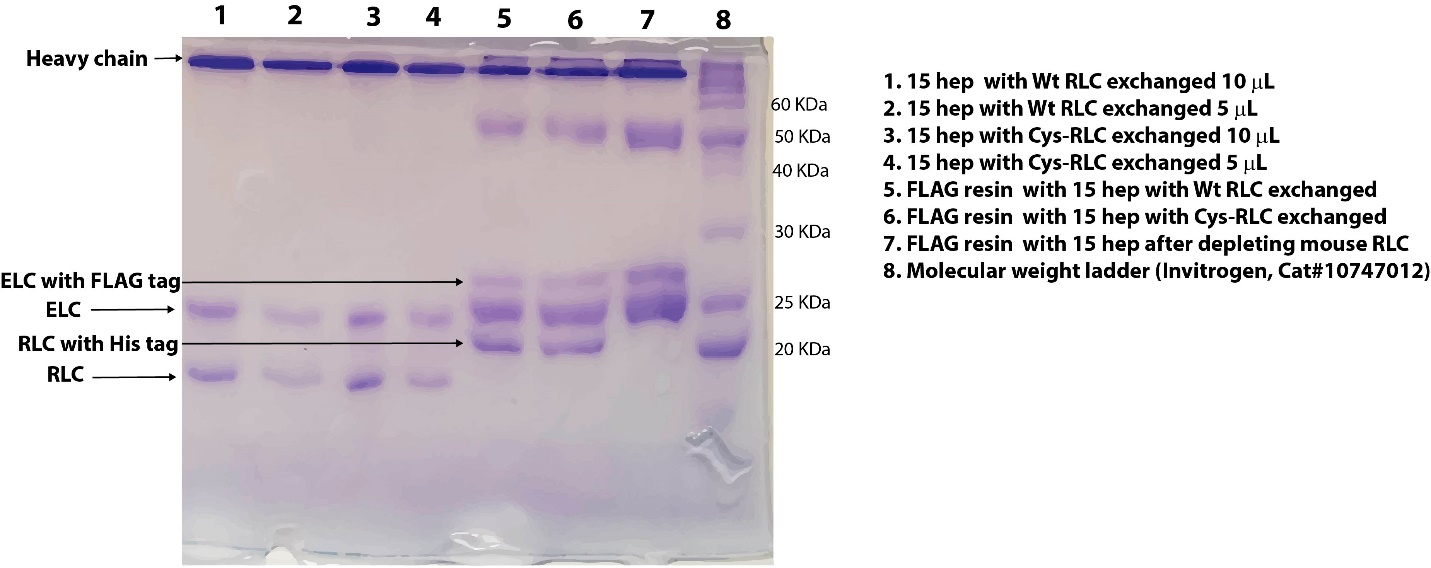

**Figure S2. Purity and extent of RLC exchange of 15-hep HMM assessed by 15% SDS PAGE. Clarified lysate from** C2C12 cells expressing 15-hep HMM heavy chain and FLAG-tagged human ELC was subjected to affinity chromatography using anti-FLAG resin followed by ion exchange chromatography using a Q Sepharose column. FLAG resin after depleting the endogenous mouse RLC was loaded in lane 7. The resin was split into two parts and one part exchanged with WT RLC and the other part exchanged with Cys-RLC (see Materials and Methods), and the samples were loaded in lanes 5 and 6 respectively. The bands at molecular weight 50 and 25 KDa in lanes 5-7 correspond to heavy chain and light chain of anti-FLAG antibody of the resin. The protein was eluted off the FLAG resin by treating with a TEV protease and further purified by Q Sepharose chromatography which gets rid of the full-length endogenous mouse skeletal myosin heavy chain (band above 15 hep HMM heavy chain in lanes 5-7) as well as the TEV protease. The purified proteins were loaded in lanes 1-4 showing bands corresponding to the WT 15-hep HMM heavy chain and the human ventricular ELC and Cys-RLC.

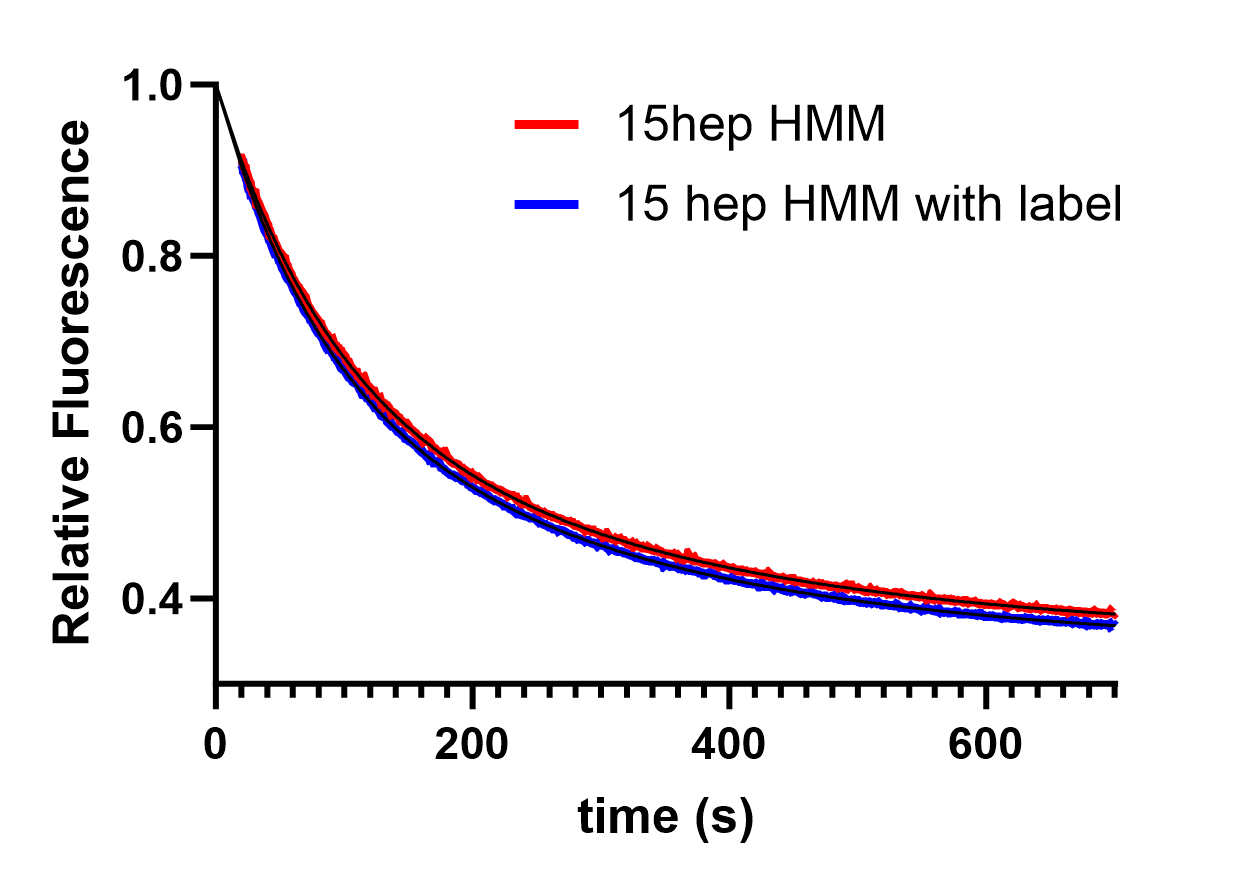

**Figure S3. Labeling of the Cys-RLC does not alter single nucleotide turnover kinetics of WT β-cardiac 15-hep HMM.** Kinetic traces are shown for WT 15-hep HMM (red) and WT 15-hep HMM with Cys-RLC labeled with AF488 (blue) at 15 mM salt. Both kinetic traces were fit to a double exponential equation, and the fits to the data are shown in black lines. The fit yielded rate constants of 0.01 s^-1^ (54% amplitude, DRX) and 0.003 s^-1^ (46% amplitude, SRX) for the unlabeled protein. The fit to the labeled protein trace yielded the same rate constants of 0.01 s^-1^ (50% amplitude, DRX) and 0.003 s^-1^ (50% amplitude, SRX).

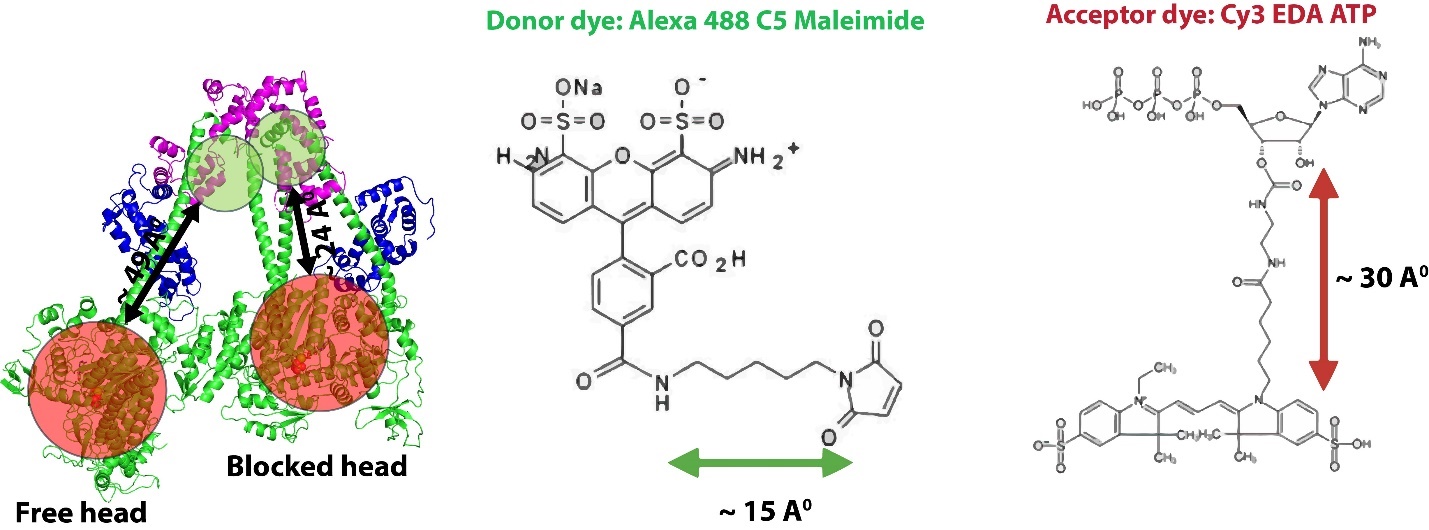

**Figure S4. Estimation of distances probed by the FRET sensor.** Schematic diagram of the IHM state of human β-cardiac 15-hep HMM showing the distances sampled by donor and acceptor fluorophores (left panel). Both the dyes used in the current FRET sensor have linkers to keep the probe away from the protein and to allow free rotation of fluorophore. The C5 linker on AF488 adds a distance of 15 Å (shown as green circle on the folded back state structure) the EDA linker on Cy3 ATP adds a distance of ~ 30 Å. Assuming the linkers sample all the available conformations, the distance between the blocked head RLC and its nucleotide binding pocket can approach as close as ~ 24 Å, while the distance for the free head can approach ~ 49 Å.
